# CTCF and R-loops are boundaries of cohesin-mediated DNA looping

**DOI:** 10.1101/2022.09.15.508177

**Authors:** Hongshan Zhang, Zhubing Shi, Edward J. Banigan, Yoori Kim, Hongtao Yu, Xiao-chen Bai, Ilya J. Finkelstein

**Affiliations:** Center for Systems and Synthetic Biology, Institute for Cellular and Molecular Biology, Department of Molecular Biosciences, University of Texas-Austin, Austin, Texas 78712, USA; Westlake Laboratory of Life Sciences and Biomedicine, Hangzhou, Zhejiang 310024, China; School of Life Sciences, Westlake University, Hangzhou, Zhejiang 310024, China; Department of Pharmacology, University of Texas Southwestern Medical Center, Dallas, Texas 75390, USA; Department of Physics and Institute for Medical Engineering and Science, Massachusetts Institute of Technology, Cambridge, MA 02139, USA; Department of New Biology, Daegu Gyeongbuk Institute of Science and Technology, Daegu 42988, Republic of Korea; Department of Biophysics, Department of Cell Biology, University of Texas Southwestern Medical Center, Dallas, Texas 75390, USA

**Keywords:** cohesin, CTCF, R-loops, single-molecule, cryo-EM

## Abstract

Cohesin and CCCTC-binding factor (CTCF) are key regulatory proteins of three-dimensional (3D) genome organization. Cohesin extrudes DNA loops that are anchored by CTCF in a polar orientation. Here, we present direct evidence that CTCF binding polarity controls cohesin-mediated DNA looping. Using single-molecule imaging of CTCF-cohesin collisions, we demonstrate that a critical N-terminal motif of CTCF blocks cohesin translocation and DNA looping. The cryo-electron microscopy structure of the intact cohesin-CTCF complex reveals that this CTCF motif ahead of zinc-fingers can only reach its binding site on the STAG1 cohesin subunit when the N-terminus of CTCF faces cohesin. Remarkably, a C-terminally oriented CTCF accelerates DNA compaction by cohesin. DNA-bound Cas9 and Cas12a ribonucleoproteins are also polar cohesin barriers, indicating that stalling is intrinsic to cohesin itself, and other proteins can substitute for CTCF in fruit flies and other eukaryotes. Finally, we show that RNA-DNA hybrids (R-loops) block cohesin-mediated DNA compaction in vitro and are enriched with cohesin subunits in vivo, likely forming TAD boundaries. Our results provide direct evidence that CTCF orientation and R-loops shape the 3D genome by directly regulating cohesin.

## Introduction

Higher eukaryotes fold their genomes into topologically associating domains (TADs) (Dekker and Mirny, 2016; Dixon et al., 2012; Nora et al., 2012; Sexton et al., 2012; Wang et al., 2016). DNA sequences within a TAD interact frequently with each other but are insulated from adjacent TADs. The cohesin complex, which is constituted of SMC1, SMC3, RAD21 and either STAG1 or STAG2, and CCCTC-binding factor (CTCF) are both enriched at TAD boundaries (Dixon et al., 2012; Guo et al., 2015; Hou et al., 2012; Merkenschlager and Nora, 2016; Nora et al., 2012; Phillips-Cremins et al., 2013; Rao et al., 2014; Sexton et al., 2012; Sofueva et al., 2013; Zuin et al., 2014). Depleting CTCF or cohesin disrupts chromosomal looping and insulation between most TADs (Haarhuis et al., 2017; Nora et al., 2017; Rao et al., 2017; Schwarzer et al., 2017; Wutz et al., 2017). CTCF defines TAD boundaries by blocking the loop extrusion activity of cohesin via an incompletely understood mechanism (Ghirlando and Felsenfeld, 2016; Li et al., 2020). TADs are also established via CTCF-independent mechanisms, including transcription and replication activities that restrict cohesin loop extrusion (Banigan et al., 2022; Dequeker et al., 2022; Jeppsson et al., 2022; Luo et al., 2022). The mechanisms underlying cohesin regulation at these roadblocks remain unclear. Here, we explore the mechanisms of CTCF-dependent and independent cohesin arrest during loop extrusion to shape the 3D genome.

CTCF arrests cohesin in an orientation-specific manner in most higher eukaryotes (Guo et al., 2015; Rao et al., 2014; Sanborn et al., 2015; Tang et al., 2015; de Wit et al., 2015). TAD boundaries are marked by CTCF-binding sites (CBSs) in a convergent arrangement. Deleting genomic CBSs abrogates TAD boundaries (Guo et al., 2015; Sanborn et al., 2015; de Wit et al., 2015) and induces aberrant gene activation (Flavahan et al., 2016; Hnisz et al., 2016; Lupiáñez et al., 2015). An interaction between an N-terminal CTCF peptide with a YDF motif and cohesin STAG1/2 subunit is essential for polar cohesin arrest and for maintaining TAD boundaries *in vivo* (Li et al., 2020). However, the mechanism(s) regulating polar cohesin arrest remain poorly explored due to the difficulty of reconstituting these biochemical activities for structure-function studies.

Topological boundaries are also established via non-CTCF mechanisms such as replication and transcription. Notably, CTCF demarcates <10% of TADs in fruit flies in orientation independent manner (Sexton et al., 2012; Hou et al., 2012; Ulianov et al., 2016; Kaushal et al., 2021). Fly TADs are depleted in active chromatin marks and separated by regions of active chromatin (Ulianov et al., 2016). The minichromosome maintenance (MCM) complex can impede the formation of CTCF-anchored cohesin loops and TADs in a cell cycle-specific manner (Dequeker et al., 2022). Chromosome-bound RNA polymerases are also capable of acting as barriers to cohesin translocation both in human and yeast (Banigan et al., 2022; Jeppsson et al., 2022). Moreover, transcribing RNA polymerases are not stationary, but rather, translocate and relocalize cohesin, which generates characteristic patterns of spatial organization around active genes (Banigan et al., 2022). Transcription products, like RNA-DNA loops (R-loops), are also postulated to reinforce TADs (Luo et al., 2022). Whether R-loops can interact with cohesin and regulate loop extrusion is unknown. These studies all point to the intriguing possibility that not only CTCF, but also additional proteins and DNA structures organize our 3D genomes.

Here, we use a combination of single-molecule studies and cryo-electron microscopy to show that CTCF and R-loops both block cohesin-mediated loop extrusion. CTCF binding polarity controls cohesin-mediated DNA looping. Cohesin that encounters the non-permissive CTCF N-terminus is blocked from further translocation and loop extrusion. The cryo-electron microscopy structure of the intact cohesin-CTCF complex reveals that this CTCF motif ahead of zinc-fingers can only reach its binding site on the STAG1 cohesin subunit when the N-terminus of CTCF faces cohesin. Remarkably, a C-terminally oriented CTCF accelerates cohesin translocation, causing increased DNA compaction. This suggests that CTCF shapes the 3D genome even when positioned in a permissive orientation relative to cohesin. DNA-bound Cas9 and Cas12a ribonucleoproteins are also polar cohesin barriers, indicating that cohesin stalling is intrinsic to this DNA motor and may be triggered by diverse proteins and/or DNA structures. Finally, we show that RNA-DNA hybrids (R-loops) are enriched with cohesin subunits *in vivo*. R-loops form insulating boundaries in the absence of CTCF and efficiently block cohesin-mediated DNA compaction *in vitro*. These results provide the first direct evidence that CTCF orientation and R-loops shape the 3D genome by directly regulating cohesin.

## Results

### CTCF is a polar barrier to cohesin translocation on U-shaped DNA

We directly visualized cohesin-mediated looping and compaction of DNA bound with CTCF (**Figure 1**). CTCF assembles into clusters of 2-8 molecules on CBSs (Gu et al., 2020; Hansen et al., 2017). We reconstituted this arrangement by inserting four co-directional CBSs into a 48.5 kb DNA substrate (**Figure 1A** and **Supplemental Methods**) (Davidson et al., 2016; Kim et al., 2017). These CTCF motifs position the CTCF N-terminus towards the right side of the DNA substrate, termed *cosR*. Full-length CTCF purified with a C-terminal MBP-Flag tag forms a stable complex with dsDNA (**Figures S1A-B**). We fluorescently labeled CTCF with Alexa488-conjugated anti-Flag antibodies. CTCF binding was visualized on aligned arrays of DNA molecules suspended above a lipid bilayer surface via total internal reflection fluorescence microscopy (**Figures 1B-C, and Movie S1**) (Kim et al., 2019). Turning buffer flow off retracted both DNA and CTCF to the barrier, confirming that CTCF is bound to the DNA (**Figure 1C**). Nearly all CTCF molecules on DNA are bound to the CBSs (**Figures 1C-D**). The half-life of CTCF bound on the CBSs is 670 ± 60 s (t_½_ ± 95% CI; N=31), which is ∼5.2-fold longer than its half-life on non-specific DNA sites (**Figure S1C**). Fewer than four CBSs significantly reduced CTCF occupancy relative to non-specific DNA binding (**Figure S1D**). We estimate that the four CBSs bind 2 ± 1 (mean ± SD; N=233) CTCF molecules, as indicated by the CTCF fluorescent intensity at the CBSs relative to the CTCF on non-specific DNA (**Figure S1E**).

**Figure 1.**
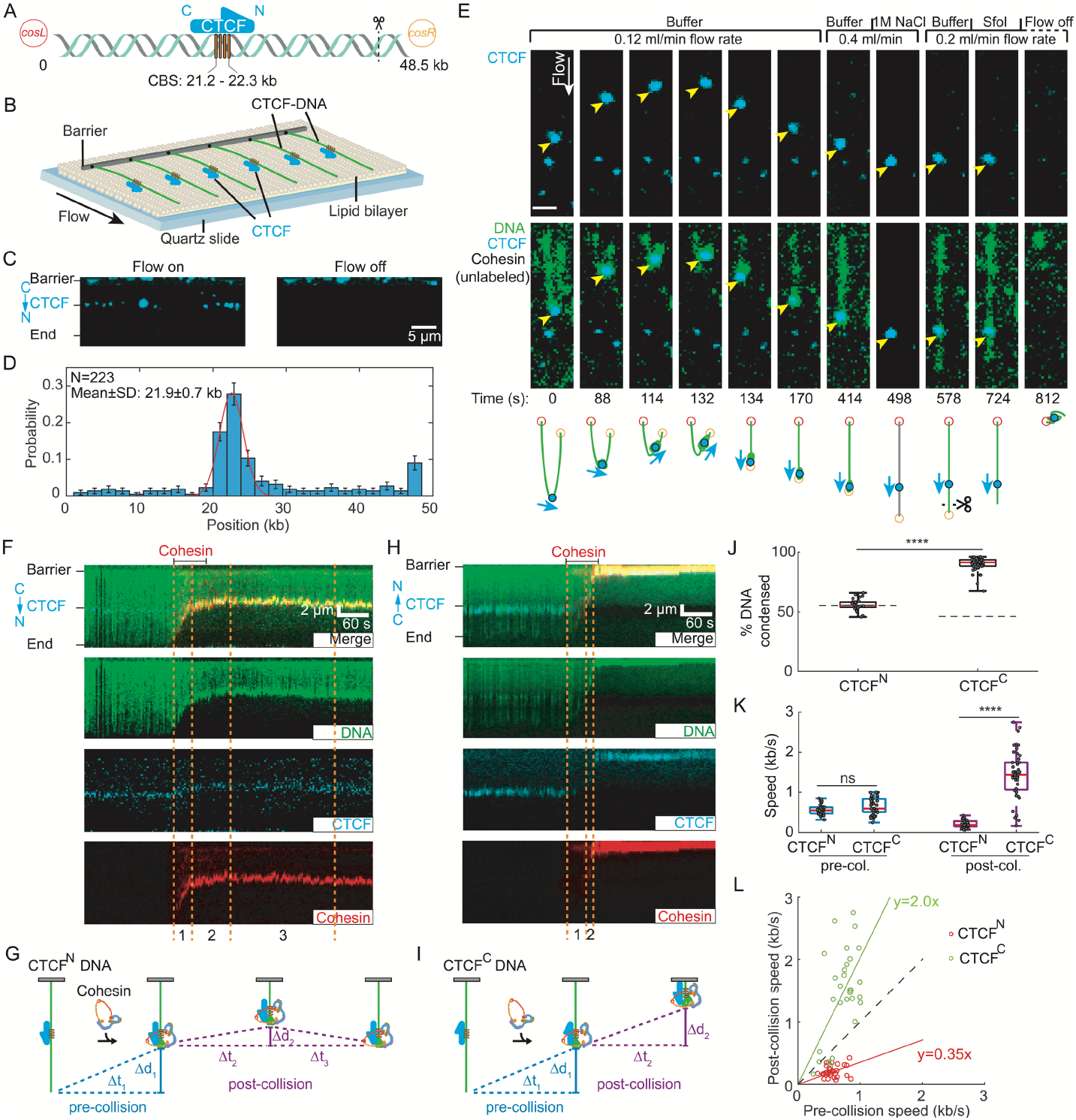
CTCF is a polar boundary for cohesin translocation. (A) Schematic of the DNA substrate. The location of the four CTCF binding sites (CBS) and the orientation of CTCF are shown in yellow boxes and a blue arrow, respectively. The black dashed line indicates the cutting site of restriction enzyme SfoI on DNA. (B) An illustration of the DNA curtain assay where the *cosL* DNA end is anchored to the flowcell surface. (C) Left: Image showing Alexa488-labeled CTCF binding to the DNA substrate. Right: Turning off buffer flow retracts the DNA and CTCF to the barrier, confirming that CTCF is bound to the DNA. (D) CTCF binding distribution on the DNA substrate. Red line: Gaussian fit. (E) Real-time visualization of CTCF stopping cohesin on U-shaped DNA. Both DNA ends are tethered to the flowcell surface. DNA is visualized with SYTOX Orange (green) and CTCF is labeled with an Alexa488-conjugated antibody (blue). Upon cohesin injection, the DNA segment between the CTCF and right tether is compacted. At 134 s, the right tether detaches from the surface, causing the left DNA segment to extend by the buffer flow. A high-salt (1M NaCl) wash at 498 s disrupts the looped DNA and washes out the SYTOX Orange stain. The DNA is re-stained by re-injecting imaging buffer. To identify the *cosR* end, we inject the restriction enzyme SfoI, which cleaves near *cosR* at 724 s. Yellow arrows show the positions of CTCF. Scale bar: 3 µm. (F) Representative 3-color kymograph showing that CTCF (labeled with Alexa488) arrests cohesin (labeled with Alexa647) in the non-permissive (CTCF^N^) orientation. Dashed lines indicate the pre- and post-collision timepoints depicted in panel G. (G) A schematic of cohesin-mediated compaction on a non-permissive CTCF-containing DNA and its analysis. Pre-collision: DNA is first condensed a distance ∆d_1_ for ∆t_1_ seconds. Post-collision: the DNA is further compacted (∆d_2_) for a short time (∆t_2_). The small DNA loop generated during ∆t_2_ is eventually dissipated (∆t_3_), as seen by the CTCF/cohesin complex returning to the pre-collision position. (H) Representative 3-color kymograph showing that CTCF permits further compaction after cohesin encounters at the permissive (CTCF^C^) orientation. Dashed lines indicate the pre- and post-collision outcomes depicted in panel I. (I) A schematic of cohesin-mediated compaction on a permissive CTCF-containing DNA and its analysis for the pre-/post-collision. DNA continues to be compacted a distance ∆d_2_ for ∆t_2_ seconds after the collision. (J) Quantification of the percentage of CTCF^N^-DNA and CTCF^C^-DNA condensed by cohesin. At least 32 DNA molecules were measured for each condition. The dashed lines indicate the CTCF binding positions on DNA substrates. (K) DNA compaction speed for the pre- and post-collisions with CTCF^N^ and CTCF^C^. Boxplots indicate the median and quartiles. P-values are obtained from two-tailed t-test: ****: P < 0.0001, ns: not significant. (L) A comparison of the speed of individual cohesins before and after colliding with CTCF^N^ (red) or CTCF^C^ (green). The dashed line with a slope of 1 is included for reference.

CTCF is a polar boundary for cohesin-mediated loop extrusion (Gómez-Marín et al., 2015; Guo et al., 2015; Rao et al., 2014; Vietri Rudan et al., 2015; de Wit et al., 2015). Cohesin that encounters CTCF from its non-permissive, N-terminal side is proposed to stop extrusion. Whether encounters from the permissive, C-terminal side of CTCF can regulate cohesin is unknown. To determine how CTCF regulates cohesin, we directly observed loop extrusion on U-shaped CTCF-DNA. In these assays, both ends of the DNA substrate are biotinylated and tethered to the flowcell surface (Ganji et al., 2018; Kim et al., 2019). DNA is visualized via the intercalating dye SYTOX Orange. Alexa488-labeled CTCF is injected into the flowcell before unlabeled cohesin-NIPBL^C^ (hereinafter referred to as cohesin) in the imaging buffer (**Figures 1E, S2, and Movies S2-S3**). After cohesin is added, a representative DNA molecule shows gradual compaction of its right arm, indicating cohesin-mediated DNA looping (**Figure 1E**) (Davidson et al., 2019; Kim et al., 2019). Notably, the left arm of DNA is not compacted completely, suggesting that CTCF acted as a polar boundary to arrest cohesin. The right end of the molecule detached at 134 s, resulting in the linearization of the looped DNA molecule. While the left arm of the DNA was gradually extended, the right arm of the DNA stayed looped. This confirms cohesin-mediated looping of the right arm of the U-shaped DNA.

Since both ends of the U-shaped DNA substrate are biotinylated, these molecules are tethered with a random CBS orientation relative to the direction of cohesin translocation. We used *in situ* optical restriction enzyme mapping to determine the polarity of the CBSs. We first washed off cohesin by injecting a high-salt buffer at an increased flow rate (**Figure 1E**). After this stringent wash, CTCF remained bound at the target site, but DNA loop was disrupted, and DNA was re-extended. The restriction enzyme SfoI cuts at a single site 2.8 kb away from the *cosR* end (**Figures 1A and S2A**). When injected into the flowcell, SfoI cut the DNA molecule at what was formerly the right arm, indicating that this was the *cosR* side of the DNA substrate. Thus, cohesin compacted the *cosR*-proximal DNA arm and the encounter of cohesin with the CTCF N-terminus arrested loop extrusion of the left DNA arm (blue arrow in **Figure 1E**). Consistent with the random tethering of U-shaped DNAs, we observed that 58% (N=42/72) of the molecules underwent complete compaction in both arms, as would be expected for a permissive CTCF-cohesin encounter (**Figure S3**). These results suggest that CTCF acts as a boundary for cohesin-mediated DNA looping when the N-terminus of CTCF is oriented towards cohesin.

### CTCF can either block or accelerate cohesin translocation

We next used a three-color single-tethered DNA curtain assay to directly visualize how CTCF regulates cohesin translocation (**Figure 1F**). The DNA was tethered to the flowcell via a streptavidin-biotin linkage on either the *cosL* side (termed CTCF^N^-DNA) or the *cosR* side (CTCF^C^-DNA). The DNA, CTCF, and cohesin (via its STAG1 subunit) were labeled with different fluorophores that could be simultaneously imaged. Consistent with our prior observations, cohesin loads near the free DNA end and rapidly compacts the substrate (Kim et al., 2019). Upon colliding with the N-terminal side of CTCF (CTCF^N^), cohesin slowed drastically and translocated a few kb upstream of the CBS. The CTCF-cohesin complex then returned to the CBS, likely via force-induced dissipation of the DNA loop (**Figures 1, F-G, and Movie S4**). All cohesin molecules stopped translocating after encountering CTCF^N^ (N=32) (**Figure 1J**).

Collisions of cohesin with the C-terminal side of CTCF (CTCF^C^) produced drastically different results. Collisions with CTCF^C^ accelerated cohesion and compacted the entire DNA molecule (N=51) (**Figures 1J-I, and Movie S5**). We compared cohesin translocation speeds before and after CTCF collisions in both orientations (**Figure 1K**). Before colliding with CTCF, cohesin speeds were indistinguishable in either CBS orientation (mean ± SD: 0.56 ± 0.13 kb s^-1^ for cohesin-CTCF^N^; 0.61 ± 0.21 kb s^-1^ for cohesin-CTCF^C^; N > 32 for both orientations). Non-permissive (CTCF^N^) collisions slowed cohesin to a velocity of 0.2 ± 0.09 bp s^-1^. Strikingly, permissive (CTCF^C^) collisions increased the cohesin velocity to 1.42 ± 0.70 bp s^-1^.

This trend was also observed for changes in the velocity of individual molecules: non-permissive collisions slowed cohesin ∼3-fold whereas permissive collisions accelerated it by ∼2-fold (**Figure 1L**). Therefore, CTCF can arrest cohesin translocation when its N-terminus is oriented towards cohesin. With its C-terminus facing cohesin, CTCF accelerates cohesin after collision, possibly to reinforce domain boundaries.

### Polar cohesin arrest requires the unstructured CTCF N-terminal domain

CTCF physically interacts with the STAG2-RAD21 cohesin subcomplex through its conserved N-terminal YDF motif (Li et al., 2020; Nora et al., 2020; Pugacheva et al., 2020). Cells with CTCF(Y226A/F228A) have fewer loops and weaker domain boundaries than wild type CTCF (Li et al., 2020). To determine whether this CTCF-cohesin interaction was required for blocking cohesin, we first characterized the ability of CTCF to arrest recombinant cohesin with STAG1(W337A/F347A; termed cohesin-WFA), which is deficient in binding the CTCF YDF motif (Li et al., 2020) (**Figures 2A-E**). Cohesin-WFA did not stop after colliding with either CTCF^N^ or CTCF^C^, and compacted the DNA in both orientations (N=66 and 30 for CTCF^N^ and CTCF^C^, respectively). The pre- and post-collision velocities were also indistinguishable in either orientation (**Figures 2D-E**). We next tested CTCF(Y226A/F228A; CTCF-YFA) and a truncation mutant that only includes the 11 Zn-fingers (CTCF-ZF) (**Figure 2F**). All CTCF mutants retained a high affinity for the CBS (**Figures S4A-B**). Strikingly, both CTCF-YFA (**Figures S4C-G**) and CTCF-ZF (**Figures S4H-L**) lost their functions as polar barriers of cohesin translocation. Thus, the interaction between STAG1 and the CTCF YDF motif is required for polar cohesin blockade.

**Figure 2.**
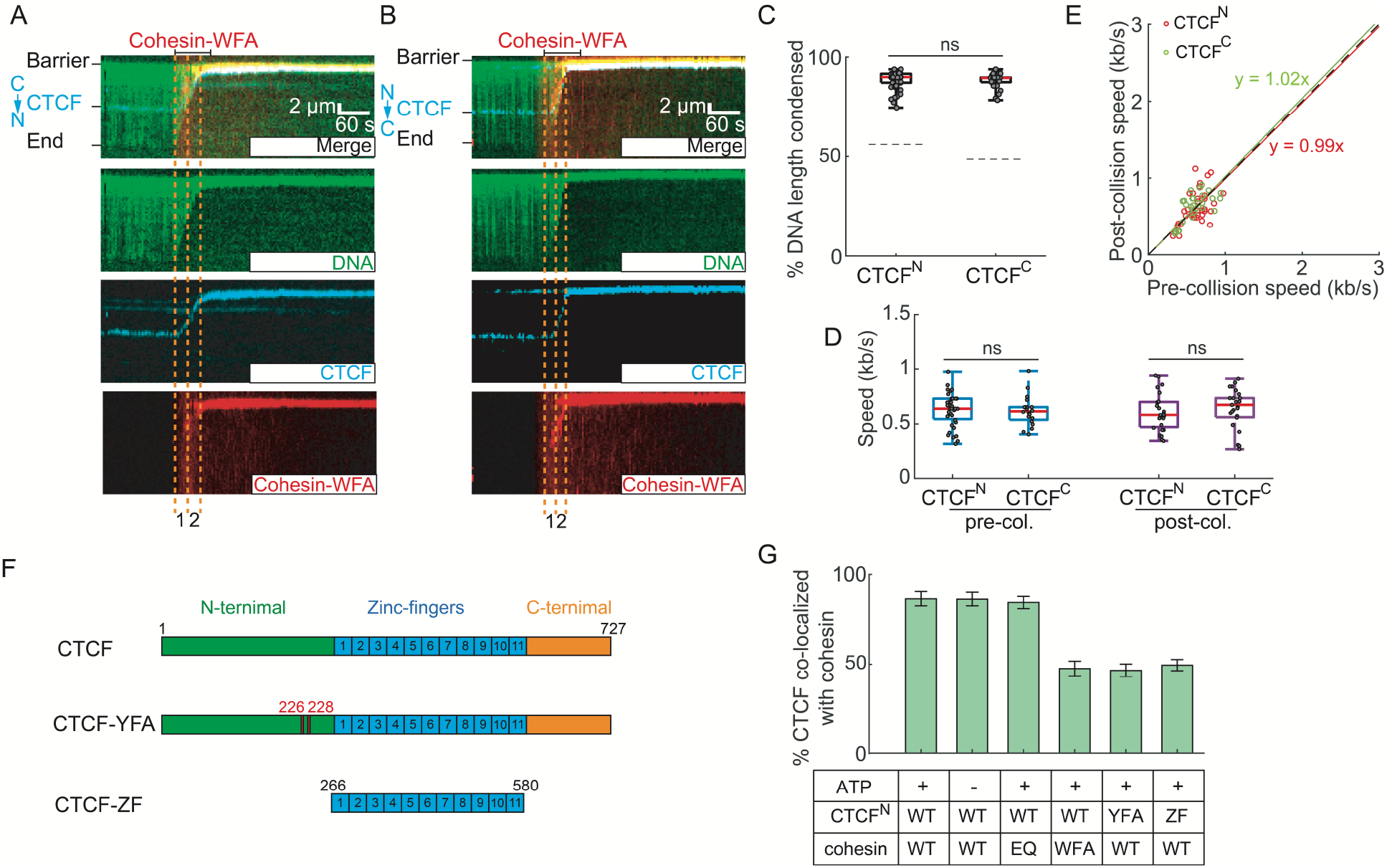
An interaction between STAG1 and the CTCF N-terminal region (NTR) are essential for polar cohesin arrest. (A-B) Representative kymographs showing that cohesin-STAG1(W337A/F347A), termed cohesin-WFA, can completely compact DNA pre-bound with (A) CTCF^N^ and (B) CTCF^C^. (C) Quantification of the CTCF^N^-DNA and CTCF^C^-DNA condensed by cohesin-WFA. The dashed lines indicate the CTCF binding positions on DNA substrates. (D) Cohesin-WFA speed pre- and post-collisions with CTCF^N^ or CTCF^C^. (E) Correlation between the speeds of individual cohesins before and after colliding with CTCF^N^ (red) or CTCF^C^ (green). The dashed line is a guide with a slope of 1. (F) Schematic of wild type CTCF, CTCF Y226A/F228A mutant (CTCF-YFA), and the zinc-finger truncation (CTCF-ZF). (G) Percent of CTCF or its mutants co-localized with cohesin variants on CTCF^N^-DNA. At least 30 DNA molecules were measured for each experiment. P-values are obtained from two-tailed t-test: ns: not significant.

CTCF-YFA and CTCF-ZF both reduced cohesin’s speed and DNA compaction in an orientation-independent manner (**Figure S4**). This observation suggests that other regions of CTCF, including the ZFs, can partially block cohesin. We thus quantified the physical interactions between cohesin and CTCF mutants. To capture potential interactions between cohesin and CTCF mutants without DNA compaction, we increased the applied laminar force to ∼0.7pN. At this force, cohesin remains bound to the DNA but cannot translocate on it (Kim et al., 2019). The vast majority of wildtype CTCF^N^ foci co-localized with cohesin (N=338/380 molecules) (**Figure 2G**). This co-localization pattern was identical without ATP and with the ATP hydrolysis-deficient cohesin SMC1A(E1157Q)/SMC3(E1144Q) mutant (cohesin-EQ). In contrast, both CTCF^N^-YFA and CTCF^N^-ZF co-localized with less than ∼50% of wildtype cohesin molecules (N=150/333 and 195/397 for CTCF^N^-YFA and CTCF^N^-ZF, respectively) (**Figure 2G**). Only 47% (N=105/224) of the wildtype CTCF^N^ foci retained cohesin-WFA. Switching the DNA orientation to CTCF^C^ resulted in a similar co-localization defect (**Figure S4M**). Thus, the CTCF-YDF motif is required for strong cohesin-CTCF binding. However, CTCF also physically interacts with cohesin via an internal region.

### Structure of the cohesin-NIPBL^C^-CTCF-DNA complex

To understand the mechanism by which CTCF blocks cohesin in an orientation-specific manner, we solved the structure of cohesin-NIPBL^C^ in complex with CTCF using cryo-electron microscopy (cryo-EM). The complex was reconstituted on a 118-base pair (bp) double-stranded DNA (dsDNA) that included a 41-bp CBS at one end. Cohesin-NIPBL^C^ remained associated with CTCF-bound dsDNA in the presence of ADP•BeF^3-^ (**Figures S5A-B**). We further stabilized this complex via mild cross-linking with BS^3^ before sucrose gradient ultracentrifugation and single-particle cryo-EM analysis.

Three-dimensional (3D) classification and refinement generated three cryo-EM maps of the complex in distinct conformations, only one of which contained one CTCF molecule bound to the cohesin-NIPBL-DNA complex (**Figures S5C-H, S6, S7 and Table S1**). The map of this conformation had an overall resolution of 6.5 Å, which allowed unambiguous rigid-body docking of the models of cohesin-NIPBL and CTCF ZFs with DNA to produce the structure of the cohesin-NIPBL-CTCF-DNA complex (Hashimoto et al., 2017; Shi et al., 2020; Yin et al., 2017) (**Figures 3A and S7A**).The CTCF N- and C-termini are predicted to be unstructured, and accordingly, are invisible in the map.

**Figure 3.**
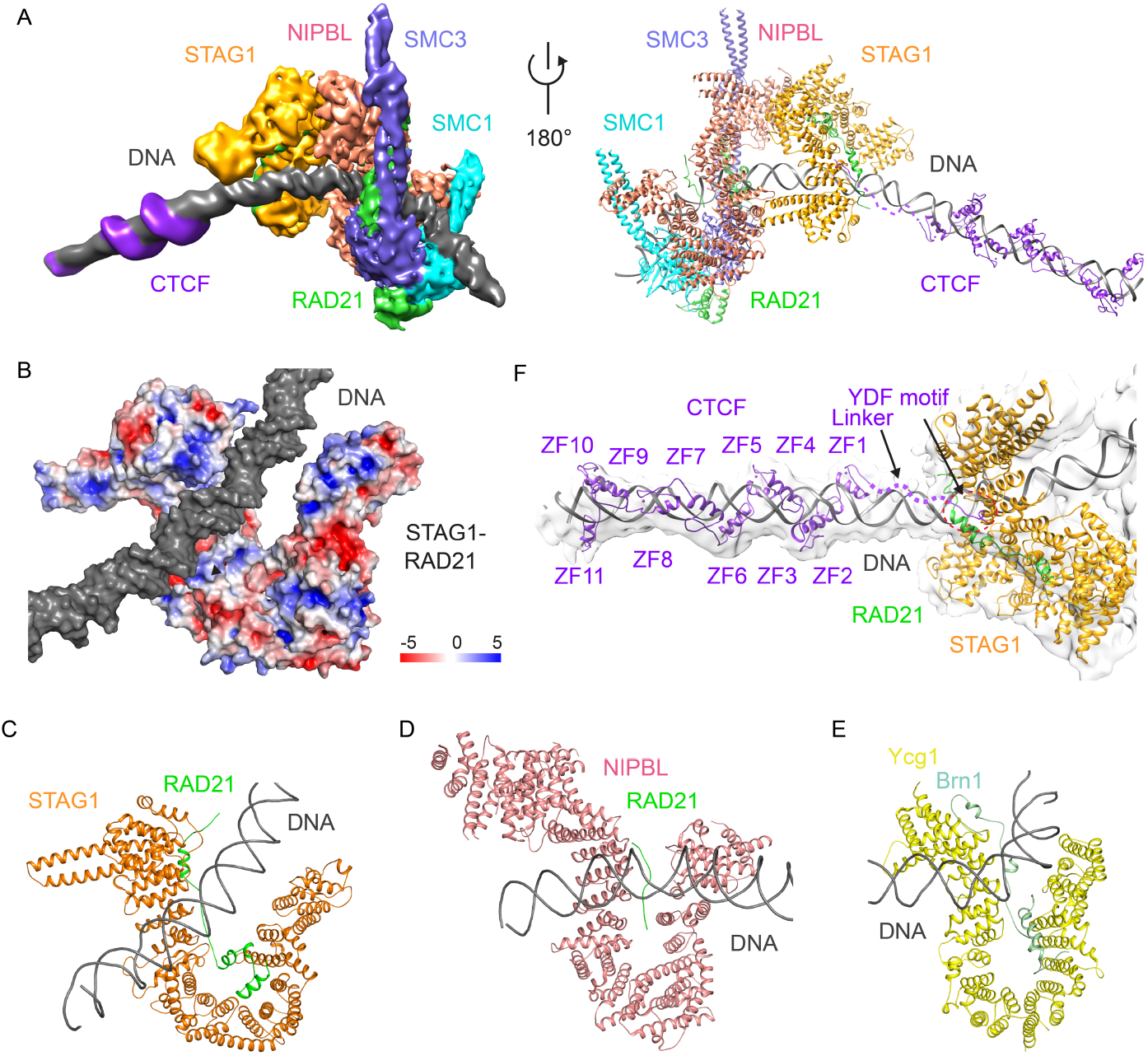
Structure of the human cohesin-NIPBL-CTCF-DNA complex. (A) Cryo-EM map (left) and model (right) of human cohesin-NIPBL-CTCF-DNA complex. DNA is captured by cohesin and NIPBL at one end and by CTCF at the other end, while its middle region contacts the top of both sides of U-shaped STAG1. (B) Surface electrostatic potential of STAG1-RAD21 subcomplex. DNA contacts positively charged regions in STAG1 and RAD21. (C-E) Structural comparison of HEAT repeat proteins STAG1 (C), NIPBL (D) and Ycg1 (E) binding to DNA duplex. (F) Locally refined map of the STAG1-CTCF-DNA subcomplex. The models of STAG1, STAG1-bound RAD21 region, DNA, and CTCF YDF motif and ZFs are shown. The CTCF linker region flanked by the YDF motif and ZFs contacts DNA.

In the complex, one end of the DNA molecule is captured by SMC1-SMC3 heterodimer and NIPBL, similar to the cohesin-NIPBL-DNA complex without CTCF (**Figure 3A**) (Collier et al., 2020; Higashi et al., 2020; Shi et al., 2020). The middle region of DNA is bound by STAG1 (**Figure 3B**). Previous studies have shown that the HEAT repeat proteins of SMC complexes, including STAG1/2 in cohesin, NIPBL, and CAP-D and CAP-G (Ycg1 in yeast) in condensin, participate in DNA binding. Unlike NIPBL that contacts DNA via its left and right arms on one side of “U” structure, STAG1 binds to DNA through both the bottom of the left arm and the tops of both arms (**Figures 3C and 3D**) (Shi et al., 2020). These DNA recognition regions in STAG1 are enriched in positively charged residues (**Figure 3B**). DNA traverses between the tops of “U”-shaped STAG1, which is similar to the DNA-binding mode by yeast condensin subunit Ycg1 (**Figure 3E**) (Kschonsak et al., 2017; Shaltiel et al., 2022). However, STAG1 possesses a wider central cleft than Ycg1, resulting in a relatively loosing binding of STAG1 to DNA, which might be an intrinsic property of cohesin. It is also possible that other unidentified factors can strength STAG1-DNA interaction at domain boundaries.

CTCF binds to the CBS at the other end of the DNA molecule, with its N-terminus pointing towards cohesin. The structure thus captures the extrusion-blocking collision complex of cohesin-CTCF. The conserved YDF motif of CTCF binds to the previously characterized site on the STAG1-RAD21 subcomplex (Li et al., 2020) (**Figure 3F**). In addition to the N-terminus, zinc finger 1 (ZF1) of CTCF contributes to cohesin positioning at CBSs, boundary insulation, and loop formation (Nishana et al., 2020; Nora et al., 2020; Pugacheva et al., 2020; Saldaña-Meyer et al., 2019). However, cohesin does not directly contact CTCF ZFs (**Figures 3A and 3F**). Thus, CTCF-ZF1 regulates cohesin via an indirect mechanism.

The YDF motif that directly binds STAG1 is conserved in CTCF proteins of various species, including *Drosophila* (**Figure S8A**). Yet, CTCF is not enriched at TAD boundaries and loop anchors in *Drosophila* (Cubeñas-Potts et al., 2017; Ramírez et al., 2018; Wang et al., 2018), suggesting that *Drosophila* CTCF cannot block cohesin. A sequence alignment of vertebrate CTCFs shows that the N-terminal region contains a patch of lysine residues close to ZF1 that may interact with DNA (**Figure S8A**). We also observed additional weak density adjacent to human CTCF ZFs on the surface of DNA in the locally refined maps (**Figure 3C**), indicating that this basic linker binds to DNA and may be important for blocking DNA compaction by cohesin. Interestingly, this basic linker is missing in *Drosophila* CTCF (**Figure S8A**), which could provide a possible explanation for the inability of *Drosophila* CTCF to stop cohesin.

### Polar arrest of cohesin by Cas9 and Cas12a ribonucleoproteins

To further probe the mechanism of cohesin arrest, we used *S. pyogenes* Cas9 as a model roadblock with a defined polarity. Nuclease-dead Cas9 (dCas9) ribonucleoprotein (RNP) was reconstituted by mixing 3xFlag-dCas9 with a single guide RNA (sgRNA). The protospacer adjacent motif (PAM) of the sgRNA faces the *cosR* end of the target DNA strand (**Figure 4A**). As expected, the dCas9 RNP labeled with a fluorescent Alexa488-anti-Flag antibody bound to its DNA target (**Figures 4B-C**) (Sternberg et al., 2014). When the DNA is tethered via its *cosL* end, cohesin collides with dCas9 from its PAM-proximal side (named dCas9^Front^) and when the DNA is tethered via its *cosR* end, cohesin encounters the PAM-distal side (dCas9^Back^; see **Figure 4A**).

**Figure 4.**
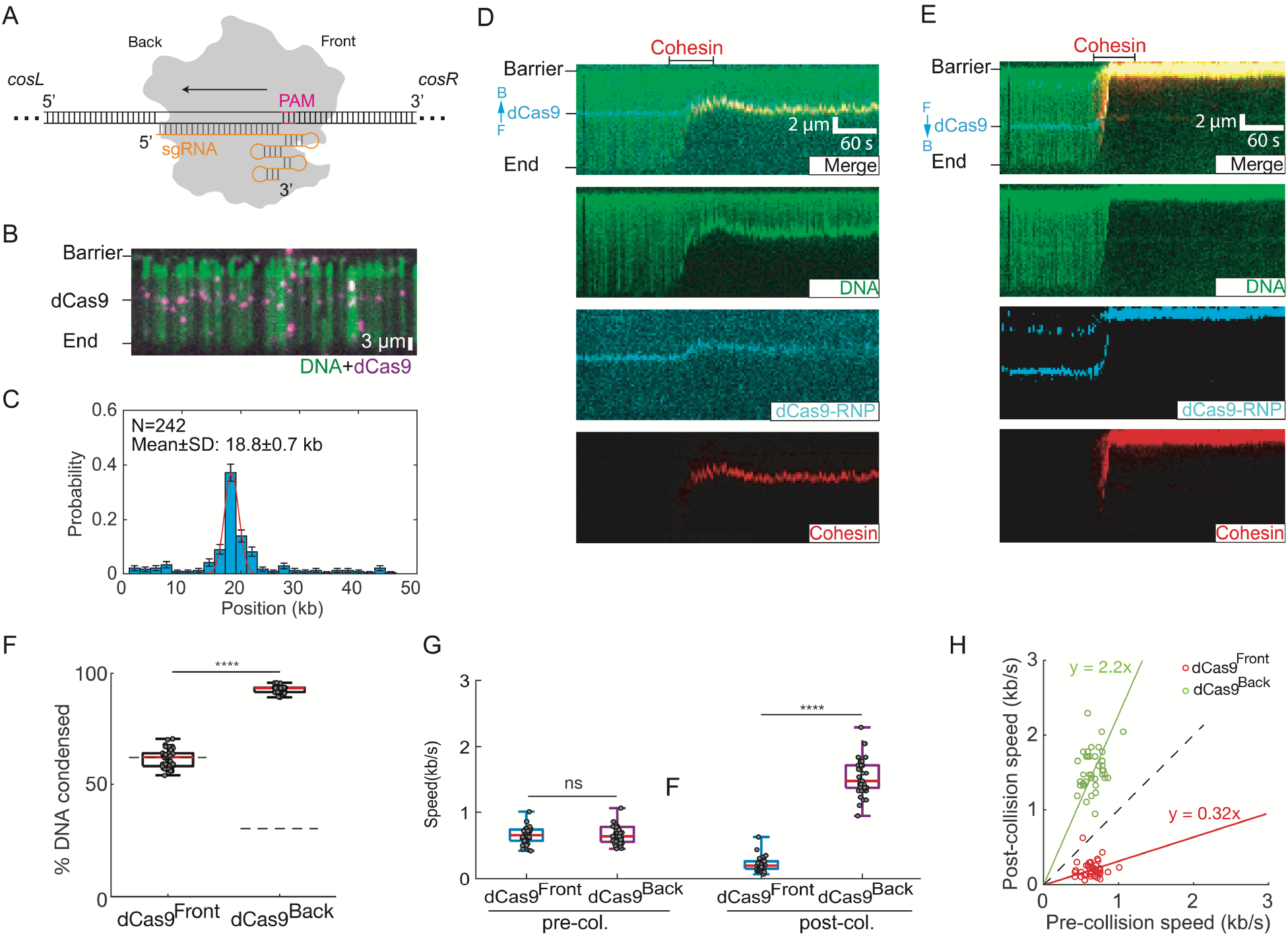
Cas9 is a polar cohesin barrier. (A) Schematic of Cas9 binding its target DNA site. sgRNA is in orange. The direction of R-loop formation is indicated with an arrow. The Cas9 protospacer adjacent motif (PAM) faces the *cosR* DNA end, termed dCas9^Front^. PAM-distal side is termed dCas9^Back^. (B) Image of Alexa488-labled dCas9 binding its target DNA. (C) Binding distribution of dCas9 on the DNA substrate. Red line: Gaussian fit. (D) Representative kymographs showing that dCas9 blocks cohesin when cohesin collides with the PAM-proximal dCas9 face (dCas9^Front^). For these experiments, the DNA is tethered via its *cosL* end. F: front, B: back. (E) When cohesin collides with the PAM-distal dCas9 face (dCas9^Back^), its post-collision speed increases. (F) Quantification of the percentage of dCas9^Front^-DNA and dCas9^Back^-DNA condensed by cohesin (N>40 for each condition). (G) Comparison of the pre- and post-collision cohesin speeds for dCas9^Front^ and dCas9^Back^. P-values are obtained from two-tailed t-test: ****: P < 0.0001, ns: not significant. (H) A scatter plot showing the relationship for individual cohesin speed before and after collision with dCas9^Front^ (red) and dCas9^Back^ (green). Dashed line is for a reference (slope = 1).

Cohesin compacts the DNA until it encounters dCas9 in either orientation. Strikingly, we observed different behaviors with dCas9^Front^ and dCas9^Back^ (**Figures 4D-E and Movies S6-S7**). dCas9^Front^ slowed cohesin ∼3-fold relative to its pre-collision speed (0.61 ± 0.12 kb/s; N=40) and eventually arrested cohesin at the collision site (**Figures 4F-G**). In contrast, collisions with dCas9^Back^ increased cohesin’s speed ∼2.2-fold (1.52 ± 0.27 kb/s; N=41) and led to nearly complete DNA compaction (**Figures 4F-H**). Thus, dCas9 recapitulates polar cohesin arrest and acceleration that we observed with CTCF (**Figure 1**). Our surprising finding that dCas9 arrests cohesin in a polar fashion explains a recent *in vitro* report that gold nanoparticle attached to dCas9-RNP only blocks ∼50% of cohesins (Pradhan et al., 2021). This partial effect is likely due to the unresolved collision polarity in that study. More importantly, our results also explain how dCas9 establishes TADs in mammalian cells (Zhang et al., 2019).

We next observed cohesin’s collisions with the nuclease inactive *Acidaminococcus* sp Cas12a (dCas12a) RNP. Cas12a and Cas9 are structurally and biochemically divergent RNA-guided nucleases. dCas12a recognizes its PAM on the 3’ side of the target DNA strand (PAM-proximal side is named dCas12a^Front^, PAM-distal side is named dCas12a^Back^; **Figure S9A**), which is opposite to dCas9 (**Figure 4A**). Fluorescently-labeled dCas12a efficiently bound its target site (**Figures S9B-C**) (Calcines-Cruz et al., 2021). dCas12a slowed and eventually stopped cohesin, but only when cohesin approached from the dCas12a^Front^ side (**Figures S9D-H**). Collisions with dCas12a^Back^ accelerated cohesin ∼1.8-fold (N=53). Together, CTCF, dCas9, and dCas12a can all arrest or accelerate cohesin, depending on the polarity of the encounter. Polar arrest is a general feature of cohesin’s translocation cycle that can be elicited by diverse roadblocks.

### R-loops act as barriers to cohesin translocation

Target-bound Cas9 and Cas12a both form an RNA:DNA hybrid (R-loop) with a displaced single-stranded DNA. R-loops also form genome-wide during transcription via hybridization of the nascent transcript with the template DNA strand. Cohesin binds RNA via its STAG1/2 subunits *in vitro* (Pan et al., 2020), and STAG1/2 proteins are enriched at R-loops in cells (Porter et al., 2021). Moreover, apoCas9 doesn’t block cohesin translocation, suggesting that R-loops may impede cohesin directly.

We generated stable R-loops *in vitro* and observed their impact on cohesin translocation (**Figure 5 and Movie S8**). R-loops were assembled via concatemerization of a plasmid encoding the mouse *Airn* gene, followed by *in vitro* transcription and RNase A treatment (**Figure S10A**) (Pan et al., 2020; Stolz et al., 2019). R-loops from the *Airn* gene are stable both in cells and *in vitro* (Pan et al., 2020; Stolz et al., 2019). The DNA concatemers varied in length from 17 to 110 kb (**Figure S10B**). Transcription did not appreciably change the distribution of DNA lengths (**Figure S10C**). We estimate 3 ± 2 R-loops per DNA molecule by fluorescently imaging these structures with the S9.6 antibody conjugated with Alexa488 (**Figures 5B and S10D**). R-loops were separated by multiples of 4 kb, as expected for a DNA substrate that is generated via multi-copy ligation of the same 4 kb-long plasmid (**Figure S10E**). These DNA molecules were biotinylated and injected into the flowcell for single-molecule imaging. About 44% of the R-loops bound cohesin, confirming a physical interaction, likely with STAG1 (**Figures 5C and S11, A to B**) (Pan et al., 2020). Cohesin did not fully compact DNA in the presence of R-loops and slowed 0.7-fold (0.42 ± 0.31 kb/s; N=37) (**Figures 5B-F**). In contrast, cohesin completely compacted non-transcribed DNA or transcribed substrates that had been digested with RNase H to remove the R-loops (**Figures 5D and S11**). As expected, R-loop substrates that were pre-treated with. RNase H did not slow cohesin (**Figure 5E**). Cohesin stalled 19 ± 8 kb after colliding with the first R-loop (N=45), indicating that a single encounter was insufficient to halt translocation. We estimate that complete cohesin arrest required 5 ± 2 R-loop collisions, on average (**Figures 5G and S11G**). A stringent 1 M NaCl wash re-extended partially looped DNA molecules, but did not disrupt the tight cohesin-R-loop interaction (**Figure 5H**). In contrast, 1M NaCl is sufficient to remove cohesin from naked DNA. We ruled out that the S9.6 antibody caused cohesin to stall by first imaging the R-loop collisions, and then labeling R-loops after the experiment was complete. We obtained similar results with both R-loop labeling approaches, indicating that R-loops indeed slow cohesin on their own (**Figures S11E-F**). Although we could not distinguish the direction of cohesin and R-loop collisions in these assays, our results indicate that multiple R-loops are sufficient to stall cohesin translocation *in vitro*, even in the absence of all transcription machinery.

**Figure 5.**
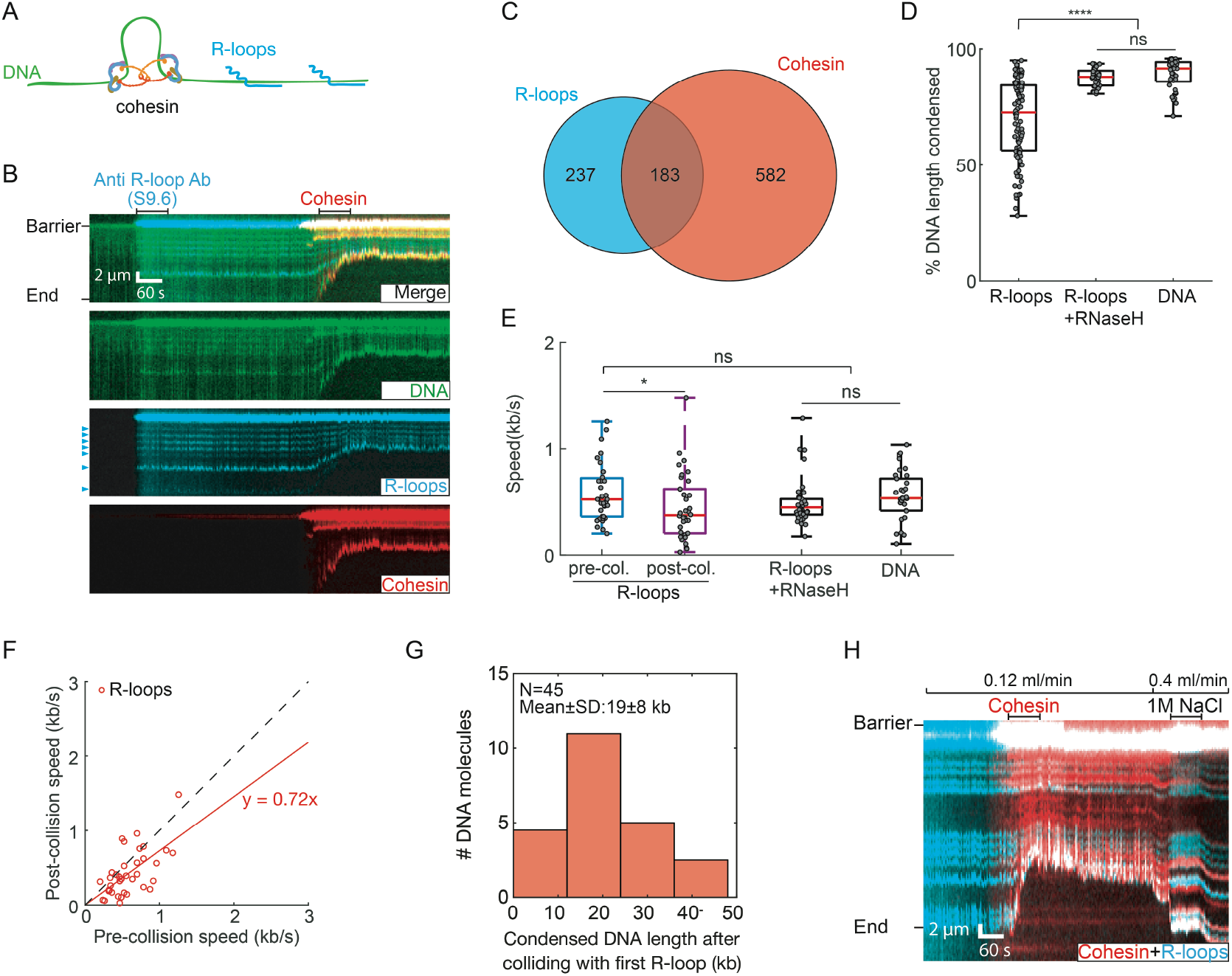
R-loops interact with cohesin and slow its translocation. (A) Schematic of cohesin translocation on the R-loops DNA substrate. (B) Representative kymographs showing cohesin colliding with R-loops. An Alexa488-conjugated S9.6 antibody is used to image the R-loops prior to cohesin injection. R-loops are indicated by arrows. (C) Venn diagram showing co-localization of R-loops and cohesin (N=110 DNA molecules). (D) R-loops significantly decrease DNA compaction, as compared with R-loops pre-treated with RNase H and non-transcribed DNA. N>38 DNA molecules for each condition. (E) After colliding with an R-loop, cohesin slows its DNA compaction. N>38 for all conditions. P-values are obtained from two-tailed t-test: *: P < 0.05, ns: not significant. (F) Individual cohesin molecules slow upon colliding their first R-loop. Dashed line is a guide with a slope of 1. (G) The counts of DNA molecules showing cohesin continues to compact DNA for ∼20 kb after colliding with the first R-loop. (H) Kymograph showing that a high salt (1 M NaCl) wash disrupts the compacted DNA. However, cohesin remains associated with the R-loop.

### R-loops are enriched for cohesin complexes and insulate genomic contacts in cells

To test whether R-loops also act as cohesin barriers in cells, we analyzed published chromatin immunoprecipitation sequencing (ChIP-seq) and DNA-RNA immunoprecipitation sequencing (DRIP-seq) datasets in mouse embryonic fibroblasts (MEFs) (Busslinger et al., 2017; Sanz et al., 2016) (**Table S4**). We analyzed chromatin localization patters for the cohesin subunits Rad21 and Stag1 as proxies for the entire cohesin complex (**Figure 6 and Table S4**). Peak overlaps were determined using the BEDTools suite with default parameters (Quinlan and Hall, 2010), and they were deemed significant by BEDTools fisher, ChIPseeker and Genomic HyperBrowser (see Methods, **Table S5**). Fifteen percent of cohesin subunit peaks overlap with R-loop peaks in WT MEFs (**Figure 6A, Table S5**). Overlaps between R-loops and Rad21 or Stag1 were nearly identical, indicating that these signals likely represent a complete cohesin complex. Knocking out the cohesin release factor Wapl did not change the cohesin and R-loop overlap genome-wide. However, the cohesin-R-loop overlap increases to ∼26% in CTCF-depleted cells and CTCF/Wapl double knock-out cells, suggesting more cohesins interact with R-loops when CTCF is ablated. Cohesin and R-loop overlap is enriched at promoter and intronic regions, consistent with the pervasive presence of R-loops between the transcription start site and the first exon-intron junction (Dumelie and Jaffrey, 2017). This enrichment is significantly enhanced in CTCF or CTCF/Wapl knock-out cell lines, indicating that R-loops provide a secondary signal for 3D genome organization (**Figure 6B**). To further confirm the significance of the overlap between cohesin and R-loop peaks, we mapped R-loop prevalence in a 5 kilobase region upstream/downstream of the cohesin-R-loop overlapped region. R-loops are prevalent around the cohesin-R-loop overlap region (**Figure 6C**), but not a few kb away from the overlaps. We also repeated this bioinformatic analysis for WT HeLa and K562 human cell lines, where both DRIP-seq and cohesin ChIP-seq datasets are publicly available (Hamperl et al., 2017; Holzmann et al., 2019; Pope et al., 2014; Sanz et al., 2016). Human cell lines also showed strong cohesin enrichment near R-loops, with the strongest enrichment at promoters and intron regions (**Figure S12, and Tables S6-S7**). We conclude that cohesin and R-loops occupy the same genomic sites, and that R-loops are additional barriers for cohesin translocation in cells.

**Figure. 6.**
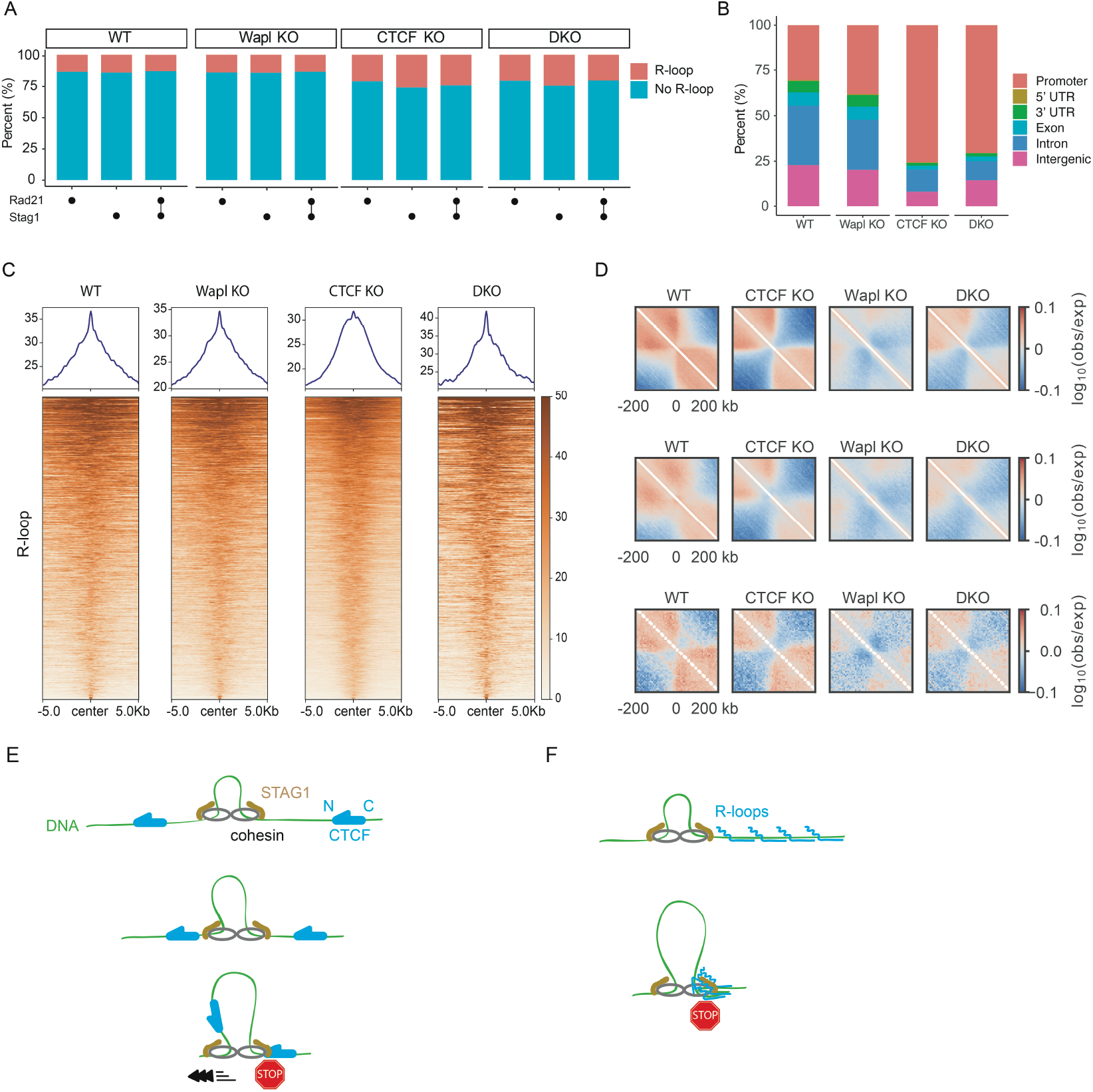
R-loops act as barriers to cohesin-mediated loop extrusion in cells. (A) Cohesin subunit Rad21 and Stag1 peak positions overlap with R-loops in WT, Wapl knockout (KO), CTCF KO, and CTCF/Wapl double KO (DKO) MEFs, as defined by ChIP-seq and DRIP-seq, respectively. Both previously published datasets were collected in mouse embryonic fibroblasts (MEFs). (B) Genomic features of overlapping regions of Rad21, Stag1 and R-loop peaks in the indicated MEFs. (C) Read density profiles and heatmaps of R-loop reads across overlaps of Rad21, Stag1 and R-loop in the indicated MEFs. (D) Average maps of chromosome contact enrichment (“observed-over-expected”; see Supplemental Methods) in MEFs (WT and mutants) in the vicinity of all R-loops (top; n=39,680; R-loops centered at 0 kb). To minimize effects of transcription start sites (TSSs) and RNA polymerase, we recomputed the maps excluding R-loops located within 10 kb of a transcription start site (middle; n=27,542). Intergenic R-loops (n=5,392) also generated insulation (bottom) in WT and mutants MEFs. (E) A summary of cohesin-regulation by CTCF. Cohesin is blocked by the N-terminus of CTCF through its interaction with STAG1 but increases its velocity when it encounters the C-terminus of CTCF. (F) A summary of the effect of R-loop clusters on cohesin translocation.

Next, we analyzed genomic contacts in the vicinity of R-loops using publicly available high-throughput chromosome conformation capture (***Hi****-****C***) and DRIP-seq datasets in MEFs (Banigan et al., 2022; Sanz et al., 2016). A map of chromosome contact enrichment averaged over and centered on oriented R-loops indicates that R-loops act as insulators for upstream and downstream contacts in WT MEFs (**Fig. 6D, top**). Insulation at R-loops depends on cohesin since the enrichment of genomic contacts vanishes in Smc3 knockout (Smc3 KO) cells (**Fig. S12D**). Knocking out CTCF somewhat increases the strength of the insulation, presumably because cohesin no longer accumulates at CTCF boundaries. When Wapl is depleted, R-loops are not insulating. This is likely because cohesin accumulates along the entire genome and has more time to traverse R-loops. A double CTCF and Wapl knockout (DKO) partially restores insulation, further demonstrating that R-loops can help shape the 3D genome. Both R-loops are RNA polymerase are enriched at promoters, and RNA polymerase generates insulation through its interactions with cohesin (Banigan et al., 2022), partially confounding our analysis. To avoid the possible effects from non-R-loop factors in promoter regions, we piled up Hi-C maps centered on R-loops but excluded those that are found within 10 kb of a transcription start site (TSS) (**Fig. 6D middle**). As an even more stringent analysis, we considered only intergenic regions while also excluding R-loops within 10 kb of a TSS (**Fig. 6D bottom**). These analyses confirm insulation near intergenic R-loops and R-loops away from TSSs, with a somewhat attenuated signal compared to the average over all R-loops. Taken together, the *in vitro* and *in vivo* analyses indicate that R-loops can act as barriers for cohesin translocation to shape the 3D genome.

## Discussion

Here, we provide direct evidence that CTCF is a polar barrier to cohesin-mediated DNA loop extrusion. Our structure of the cohesin-NIPBL-CTCF-DNA complex represents the extrusion-arrested state. In this state, cohesin adopts a fully folded conformation, with STAG1/2 engaging the N-terminal YDF motif of CTCF. This arrangement strongly suggests that as cohesin translocates on DNA, STAG1/2 is positioned in the front end of the complex. When cohesin approaches the N-terminus of CTCF, the YDF motif in the N-terminal region of CTCF can interact with STAG1/2, thus blocking cohesin translocation (**Figures 6E & S13A**). Our structure thus explains the molecular basis of the polar cohesin arrest by CTCF.

Cohesin translocation on DNA is accelerated when it first encounters the C-terminus of CTCF (**Figure 6F**). Such acceleration further improves cohesin’s processivity and may be important for reinforcing domain boundaries in cells. This acceleration requires the interaction between STAG1/2 and the YDF motif of CTCF. However, the extrusion-arrested structure reported here suggests that the N-terminal CTCF linker, which directly contacts DNA region ahead of ZF-binding site, may not allow the YDF motif to reach STAG1/2 when cohesin approaches from the C-terminus of CTCF (**Figures S13B-C**). We speculate that CTCF might interact with STAG1/2 in an actively extruding cohesin-NIPBL complex, and that this interaction further activates cohesin translocation. The molecule basis for the increased cohesin velocity is unclear. One possibility is that CTCF suppresses cohesin’s tendency to slip on DNA, especially at higher applied forces. By directly interacting with STAG1/2, CTCF may act as a processivity factor that prevents microscopic cohesin slipping during its translocation cycle to reinforce cohesin’s loop extrusion at high tensions in mammalian cells. Additional structural and biochemical studies are needed to fully elucidate how cohesin extrudes loops, and how CTCF reinforces this process.

Remarkably, both Cas9 and Cas12a RNPs can recapitulate polar cohesin arrest and acceleration, suggesting that CTCF is not unique in this regard. We only observed polar arrest/acceleration for the RNP but not the apo-Cas9/Cas12a complexes, suggesting that the protein-generated R-loop is important for this activity. Although cohesin has not evolved to see such RNPs *in vivo*, this result hints that *S. pombe, S. cerevisiae, C. elegans*, and *A. thaliana* may not need a CTCF homolog to organize their genomes. These organisms can form distinct chromatin domains reminiscent of TADs seen in humans (Rowley and Corces, 2016). Moreover, *Drosophila* CTCF performs fundamentally different functions from the human homolog, and its chromosome contact domains can form without stabilized point-to-point border interactions between CTCF sites (Rowley et al., 2017). Thus, cohesin must recognize additional CTCF-independent signals to form TADs, and these may include unidentified DNA-binding proteins or nucleic acid structures.

Even in human cells, some TAD boundary elements are not CTCF-dependent, suggesting that additional principles can also establish chromosome contact domains (Rao et al., 2014). Here, we show that R-loops can arrest cohesin, and that cohesin is enriched at R-loops *in vivo*. Our *in vivo* co-localization and Hi-C analyses are correlative and cannot rule out an indirect mechanism for R-loop associated cohesin and contact enrichment, possibly via RNA polymerase-mediated insulation at these sites. Future experiments will be required to directly test this hypothesis. Additional evidence for the importance of R-loops includes the formation of fine-scale chromatin loops connecting the promoter, the enhancer, and downstream exon regions soon after induction of transcription (Pezone et al., 2019). Interestingly, RNase H1 destroys these formed loops and eliminates cohesin binding to these sites, suggesting that these chromatin loops depend on R-loops and cohesin. Moreover, accumulation of RNA-DNA hybrids flanking CBSs decreases CTCF binding to CBSs in DIS3-deficient B cells and disorganizes cohesin localization, negatively impacting the integrity of the TAD containing the immunoglobulin heavy chain (Igh) locus (Laffleur et al., 2021). Active transcription also limits cohesin-mediated loop extrusion during RAG scanning (Zhang et al., 2019). We conclude that R-loops can arrest cohesin-catalyzed DNA looping both *in vitro* and *in vivo* with broad implications for the roles of R-loops and other roadblocks in shaping 3D genome organization in cells.

## Author Contributions and Notes

H.Z. designed and performed all single-molecule assays, analyzed single-molecule data as well as DRIP/ChIP-seq data. Z.S. performed protein purification. Z.S. and X.-c.B. performed cryo-EM study. Y.K. contributed to project design and protein purification. E.J.B. performed the Hi-C analysis. H.Y., X.-c.B. and I.J.F. co-supervised the project. All authors contributed to the writing of the manuscript.

## Acknowledgments

We thank the members of Finkelstein, Yu and Bai labs for useful discussion, and the staff of the Cryo-Electron Microscopy Facility at University of Texas Southwestern Medical Center (UTSW) for technical support.

## Data and materials availability

All data are available in the manuscript or the supplementary material. The cryo-EM map and coordinate have been deposited in the Electron Microscopy Data Bank under accession code EMD-32252 and the Protein Data Bank with ID 7W1M.

