## Supplement for "CTCF and R-loops are boundaries of cohesin-mediated DNA looping"

### SUPPLEMENTARY FIGURES

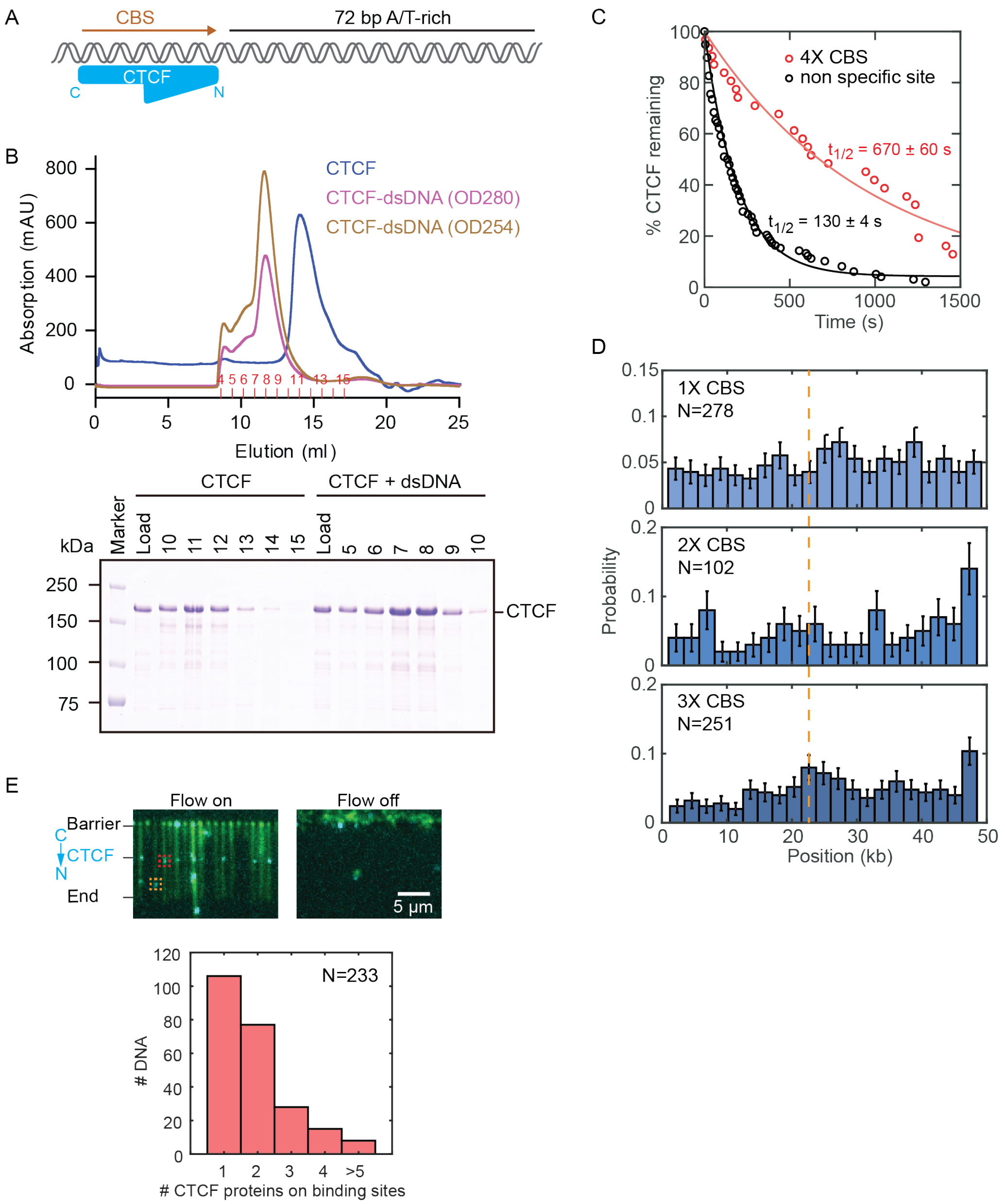


**Figure S1. Purification and biochemical characterization of the CTCF-DNA complex. Related to Figure 1.**

(A) Illustration of CTCF bound to the CTCF binding site (CBS).

(B) Size exclusion chromatography filtration profiles and corresponding SDS-PAGE analysis of CTCF and CTCF-DNA complexes.

(C) CTCF lifetimes on the 4X CBS DNA substrate (N=31) and on non-specific DNA sites (N=98). Solid lines are single exponential fits to the data. The half-life of each curve is indicated.

(D) CTCF binding distribution on DNA with single, double, and triple CBS. See Figure 1 for a binding distribution on the 4X CBS DNA substrate.

(E) We estimated the number of CTCF proteins on the 4x CBS DNA substrate by comparing the fluorescence intensity of the CTCF signal on the CBS (red box, N=233) to the average fluorescence intensity of CTCF patches on non-specific sites (orange box, N=237). Flow on/off indicates the specific and non-specific CTCF binding on DNA.

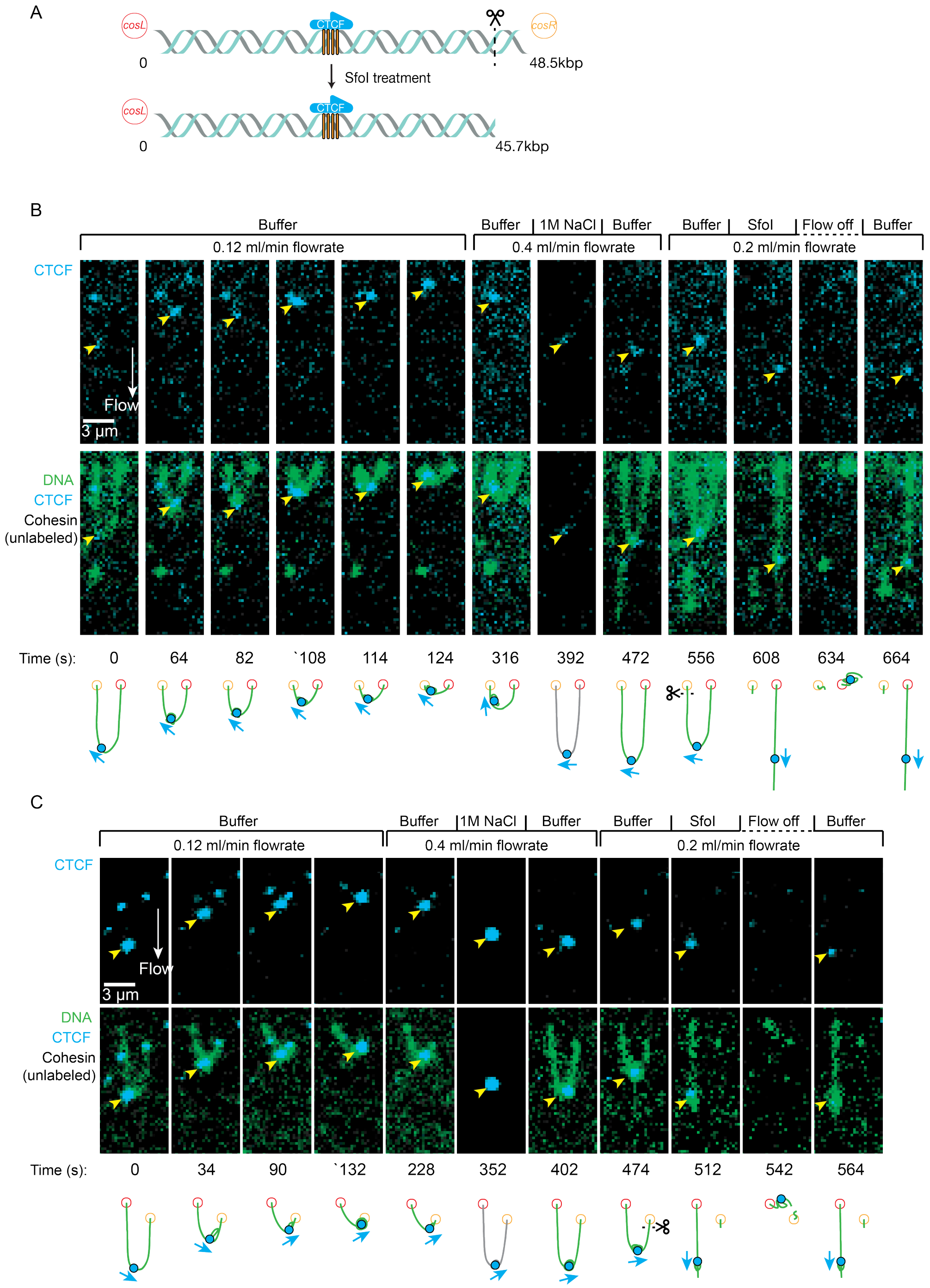


**Figure S2. Cohesin compacts the segment between CTCF and *cosR* on U-shaped DNA. Related to Figure 1.**

(A) The orientation of DNA is determined via optical restriction enzyme mapping with SfoI, which cleaves 1.2 kb away from *cosR*.

(B-C) Two examples of U-shaped DNA showing cohesin compacts the DNA segment on the N-terminal side of CTCF. Images are single frames taken at the indicated time of a single molecule movie.

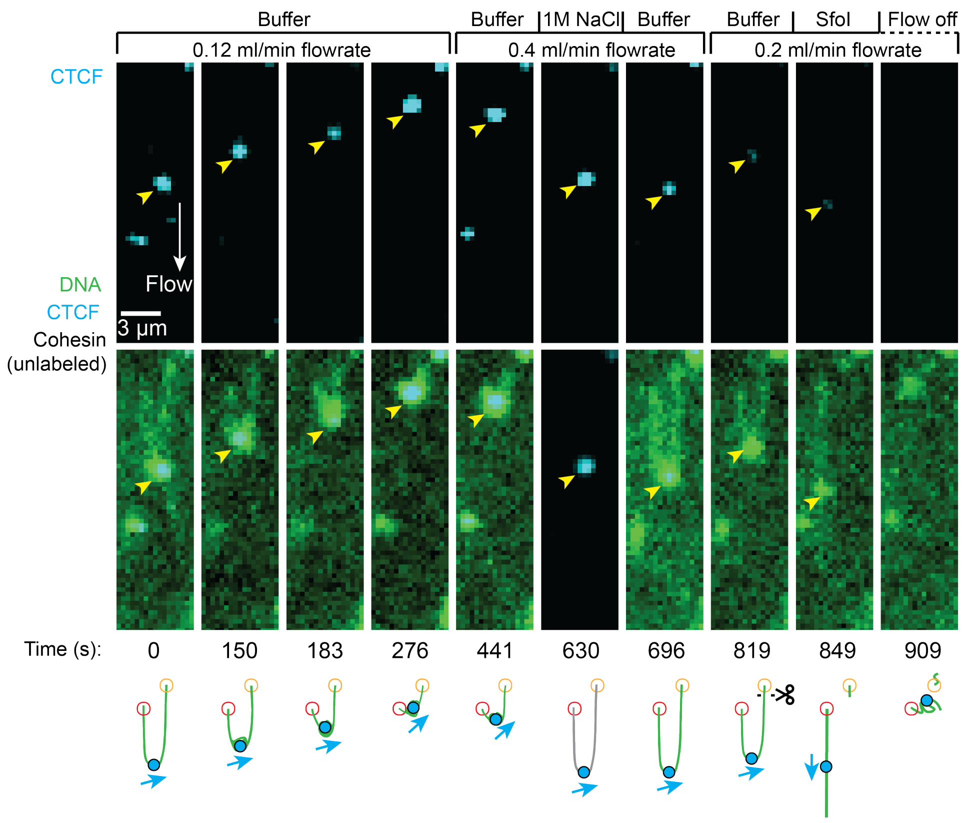


**Figure S3. Examples of cohesin compacting both segments of U-shaped DNA when it encounters CTCF$^{C}$. Related to Figure 1.**

Both DNA ends are tethered to the flowcell surface. DNA is visualized with SYTOX Orange (green) and CTCF is labeled with an Alexa488-conjugated antibody (blue). DNA is compacted upon cohesin injection, which is disrupted by 1 M NaCl. Restriction enzyme mapping with SfoI, which cleaves near *cosR,* is used to identify the orientation of CTCF. Yellow arrows show the positions of CTCF.

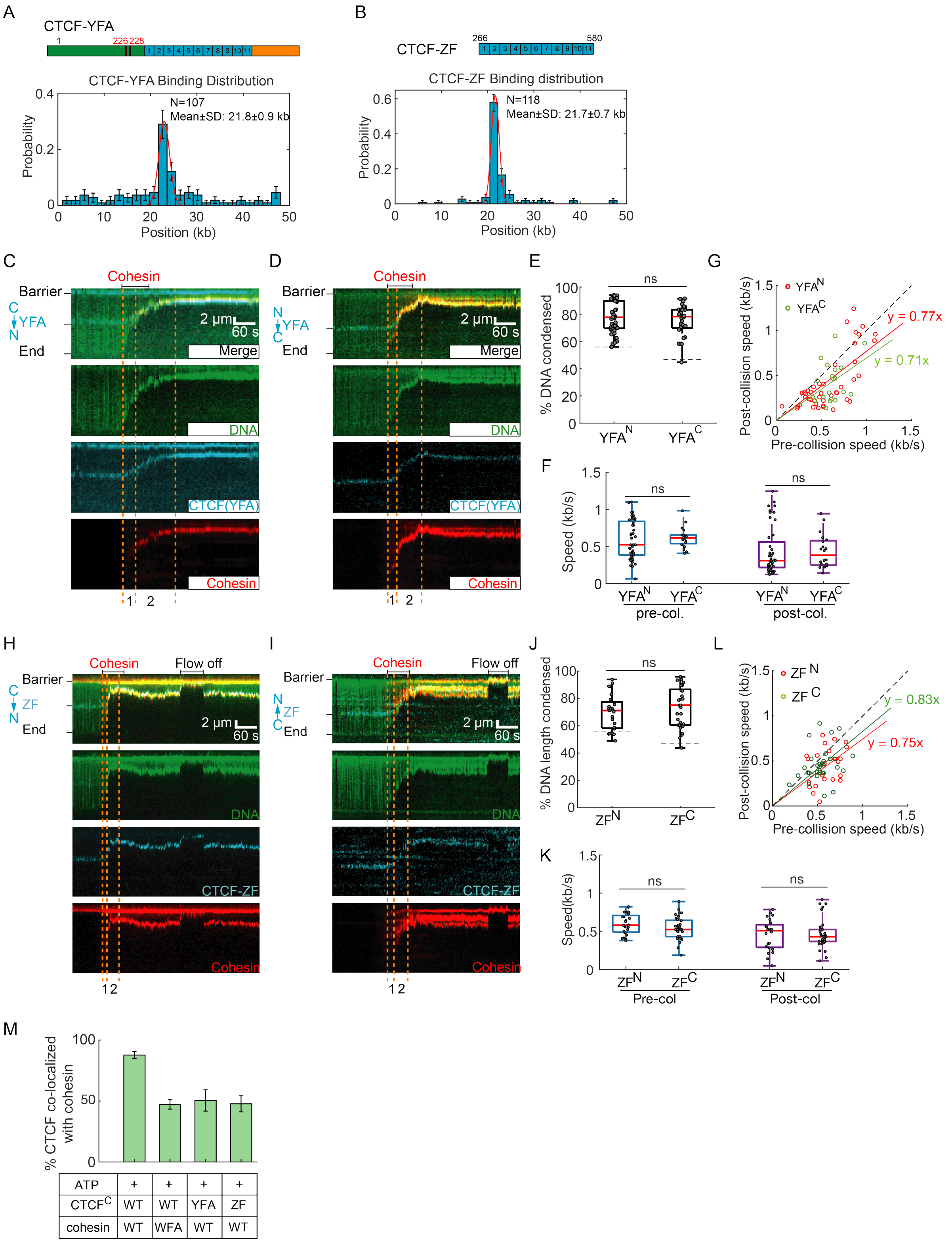


**Figure S4. Characterization of CTCF mutants and their collision with cohesin. Related to Figure 2.**

(A) CTCF-YFA and (B) CTCF-ZF binding distributions on DNA. Red curves: Gaussian fit. The center and S.D. of the fit are indicated in the histograms.

(C,D) Representative kymographs showing that cohesin compacts DNA after encountering $CTCF^N$-YFA and $CTCF^C$-YFA, respectively.
(E) Quantification of the compaction of DNA with $CTCF^N$-YFA (N=44) and $CTCF^C$-YFA (N=28).
(F) Cohesin speed pre- and post-collision with $CTCF^N$-YFA and $CTCF^C$-YFA. Dashed line: location of the CTCF binding sites on the DNA. Box plots indicate the median and quartiles.
(G) Pre- and post-collision speeds of individual cohesin molecules on DNA with $CTCF^N$-YFA (red) and $CTCF^C$-YFA (green). Dashed line is a slope of one to guide the eye.
(H,I) Representative kymographs showing that cohesin compacts $CTCF^N$-ZF and $CTCF^C$-ZF DNA, respectively.
(J) Quantification of the compaction of DNA with $CTCF^N$-ZF (N=24) and $CTCF^C$-ZF (N=33).
(K) The speed of cohesin translocation pre- and post-collision with $CTCF^N$-ZF and $CTCF^C$-ZF.
(L) Pre- and post-collision speeds of individual cohesin complexes on DNA with $CTCF^N$-ZF (red) or $CTCF^C$-YFA (green). Dashed line is a slope of one.
(M) The percent of CTCF variants co-localized with cohesin on $CTCF^C$ DNA (N>245 molecules for each condition from at least three flowcells).

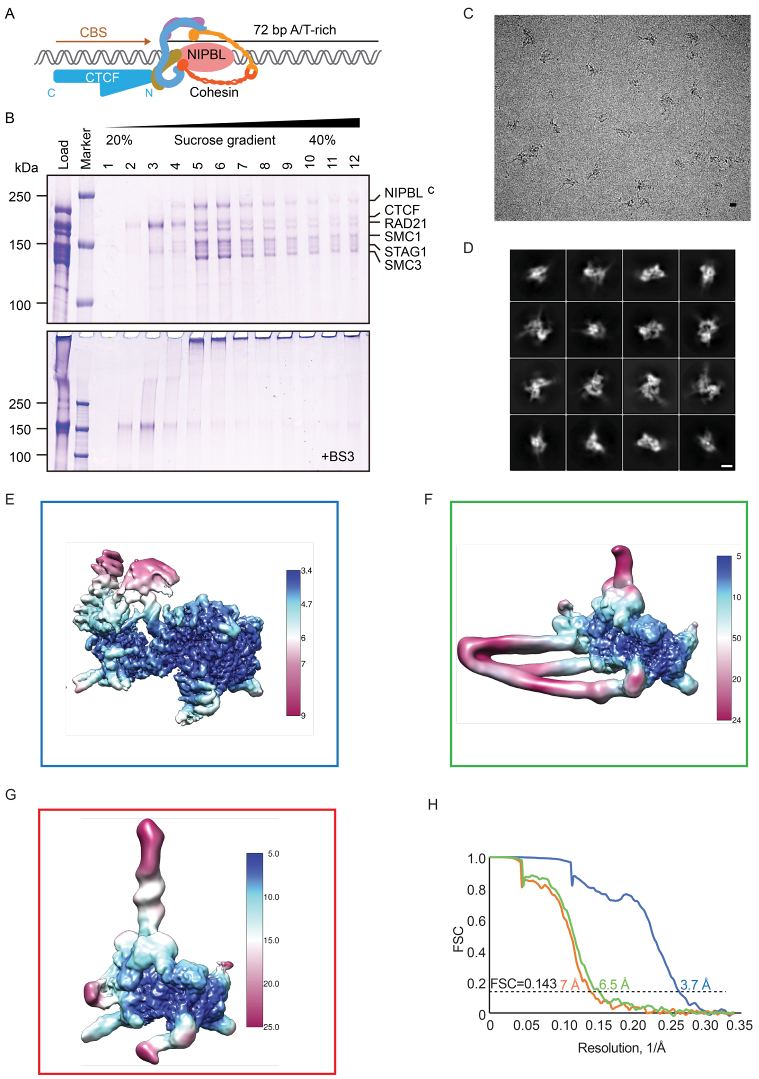


**Figure S5. Cryo-EM sample preparation and structure determination of the human cohesin-NIPBL-CTCF-DNA complex. Related to Figure 3.**

(A) An illustration of cohesin, NIPBL and CTCF bound to the DNA substrate.

(B) SDS-PAGE analysis of the cohesin-NIPBL-CTCF-DNA complex with and without mild bis(sulfosuccinimidyl)suberate (BS3) crosslinking in the presence of ADP and $BeF^{3-}$ after sucrose gradient ultracentrifugation.

(C) A representative micrograph of human cohesin-NIPBL-CTCF-DNA complex. Bar: 100 Å. (D) 2D class averages were selected for 3D reconstruction. Bar: 100 Å.

(E-G) Local resolution of the cryo-EM maps of the cohesin-NIPBL-DNA complex (E), the cohesin-NIPBL-DNA complex with folded coiled-coils (F), and the cohesin-NIPBL-CTCF-DNA complex (G). The box colors correspond to the curves in (H).

(H) Fourier shell correlation (FSC) curves of the cohesin-NIPBL-DNA complex (blue curve), the cohesin-NIPBL-DNA complex in folded state (orange curve), and the cohesin-NIPBL-CTCF-DNA complex (green curve).

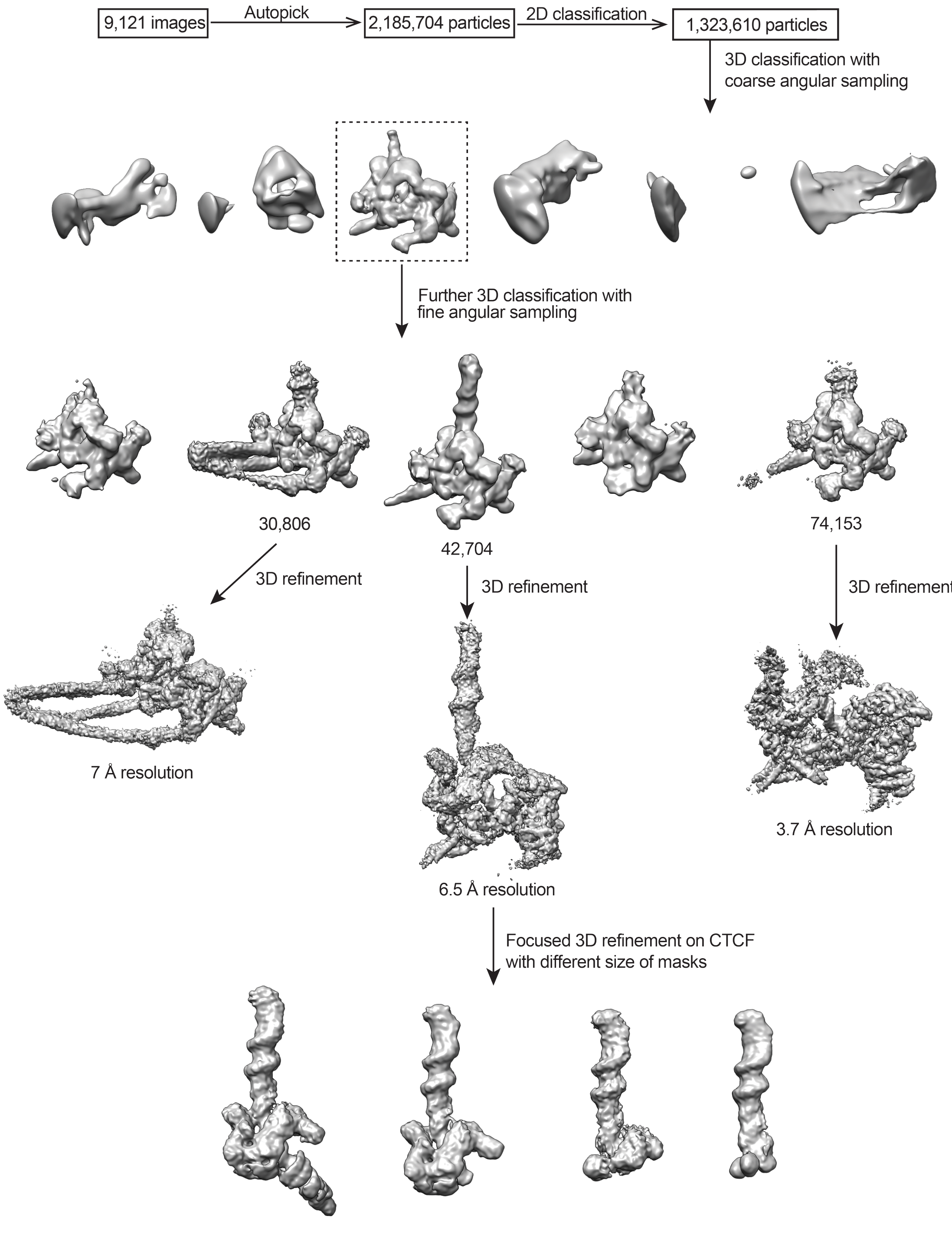


**Figure S6. Cryo-EM data processing workflow. Related to Figure 3.**
More than 2.1 million particles were picked from 9,121 images, and then were classified by 2D and 3D classification and refinement, generating three cryo-EM maps of the complex in distinct conformations.

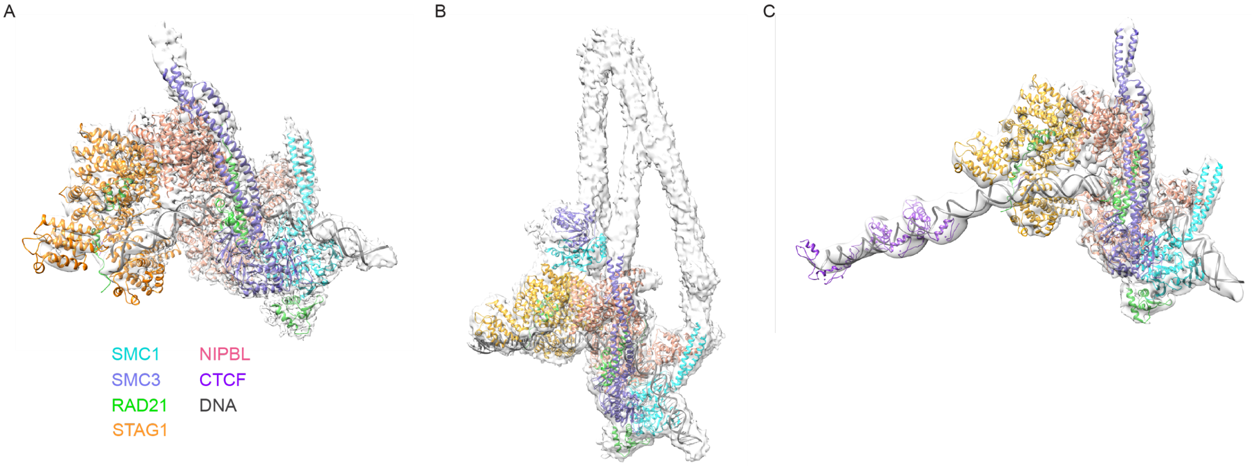


**Figure S7. Models of cohesin, NIPBL and CTCF were docked to cryo-EM maps. Related to Figure 3.**

(A) The cohesin-NIPBL-DNA complex at 3.7 Å resolution.

(B) The cohesin-NIPBL-DNA complex in folded state at 7.0 Å resolution.

(C) The cohesin-NIPBL-CTCF-DNA complex at 6.5 Å resolution. Maps and models are shown as transparent surfaces and cartoons, respectively.

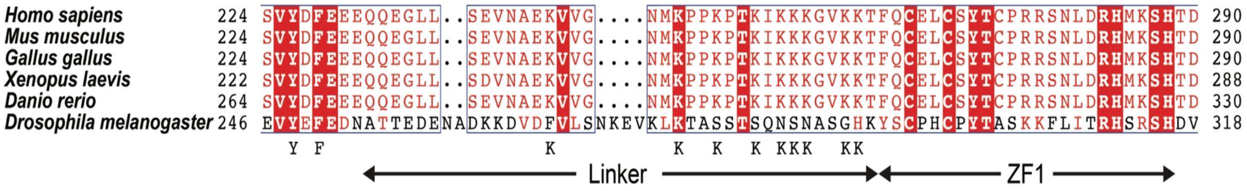


**Figure S8. Sequence alignment of CTCF from different species. Related to Figure 3.**
The linker region between the YDF motif and zinc fingers is highly conserved from zebrafish to human but is divergent in fruit fly. The conserved YXF motif and conserved lysine residues are indicated below the diagram.

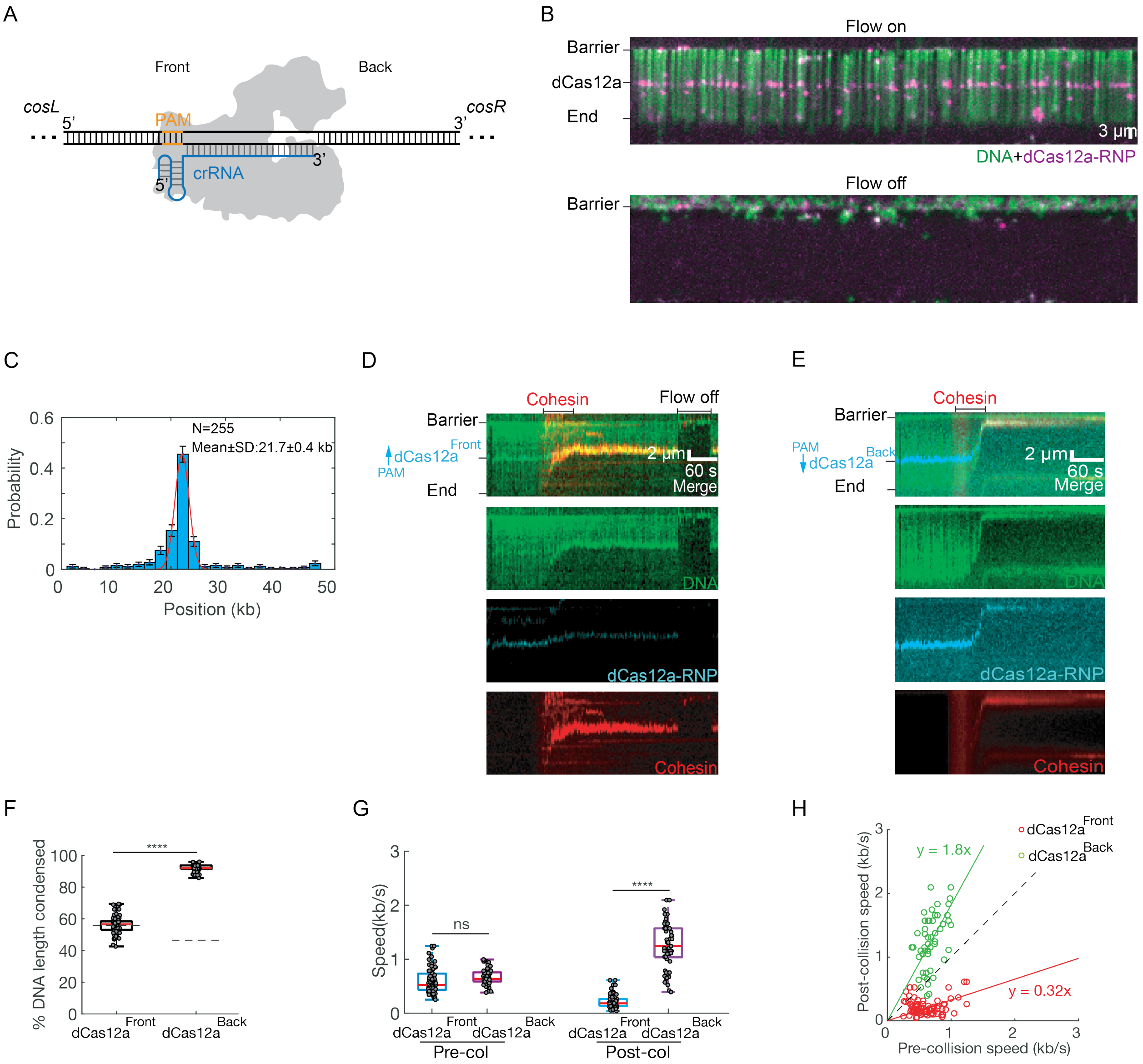


**Figure S9. Nuclease dead (d) Cas12a is a polar barrier to cohesin translocation. Related to Figure 4.**

(A) Illustration of dCas12a binding on its target site. Orange: protospacer adjacent motif (PAM).

(B) Image of Alexa488-labled dCas12a on DNA with buffer flow on (top) and off (bottom). The position of the DNA target is indicated on the left.

(C) Binding distribution of dCas12a on DNA. Red curve: Gaussian fit.

(D-E) Representative kymographs showing that $dCas12a^{Front}$ blocks cohesin (D), whereas $dCas12a^{Back}$ accelerates cohesin translocation (E).

(F) Quantification of the condensed DNA in both $dCas12a^{Front}$ and $dCas12a^{Back}$ orientations (N>45 DNA molecules for each condition). Dashed line: dCas12a position on the DNA.

(G) Pre- and post-collision speeds for $dCas12a^{Front}$ and $dCas12a^{Back}$.

(H) Correlation between the pre- and post-collision speeds of individual cohesin for the dCas12a$^{Front}$ (red) and dCas12a$^{Back}$ (green) orientations.

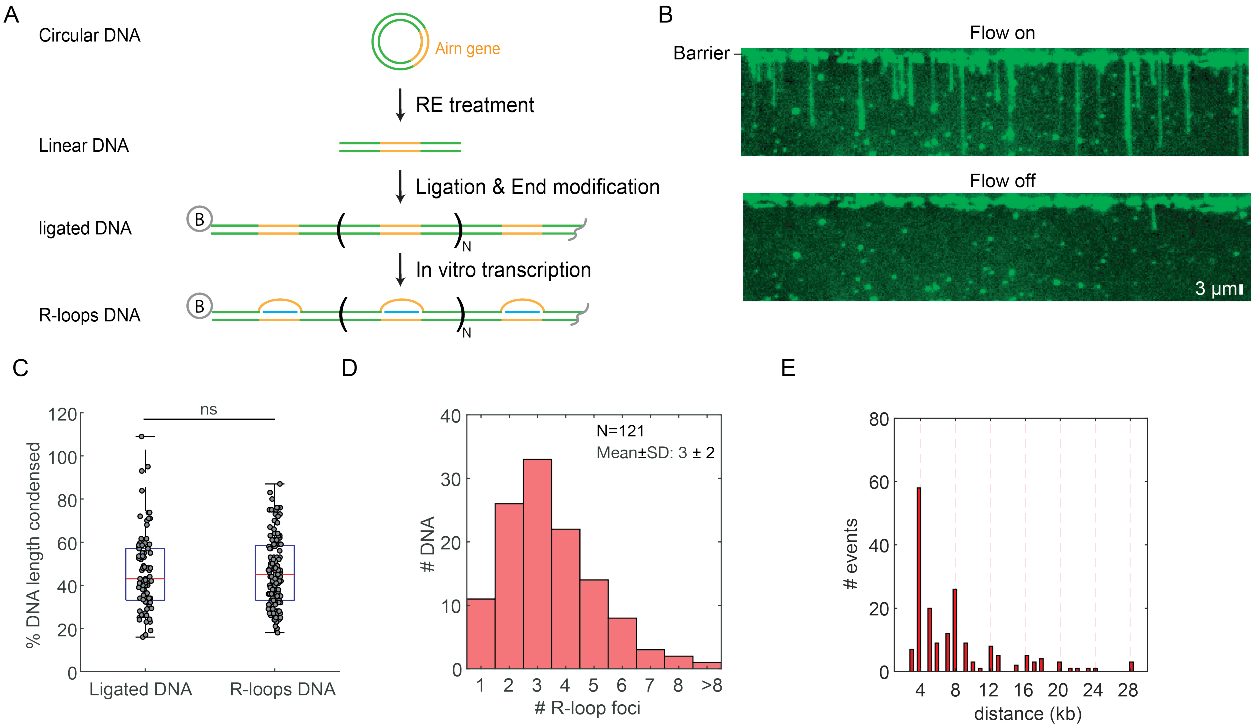


**Figure S10. Generation and characterization of the R-loops DNA substrates. Related to Figure 5.**

(A) Overview of how the R-loop DNA concatamers are generated. The *Airn* DNA sequence (orange) forms a stable R-loop when transcribed *in vitro* (Pan et al., 2020; Stolz et al., 2019). This sequence is cloned into a 4 kb plasmid. The plasmid is linearized with a restriction enzyme (RE), ligated into concatamers, and transcribed with T3 RNA polymerase. An optional RNase A treatment removes RNA that is outside the R-loop (Pan et al., 2020). B: biotin.

(B) Fluorescent image showing the distribution of DNA concatemer lengths after ligation. DNA is stained with SYTOX-Orange.

(C) The distribution of non-transcribed and transcribed ligated DNA (R-loops DNA) lengths. N>81 for both conditions.

(D) Number of R-loop foci per DNA molecule. The foci were stained with an Alexa488-conjugated S9.6 antibody.

(E) The distance between adjacent S9.6 antibody pairs (N=188). Dashed lines indicate multiples of 4 kb. At the experimental condition (0.12 mL min$^{-1}$ flowrate and 100 nM SYTOX Orange in imaging buffer).

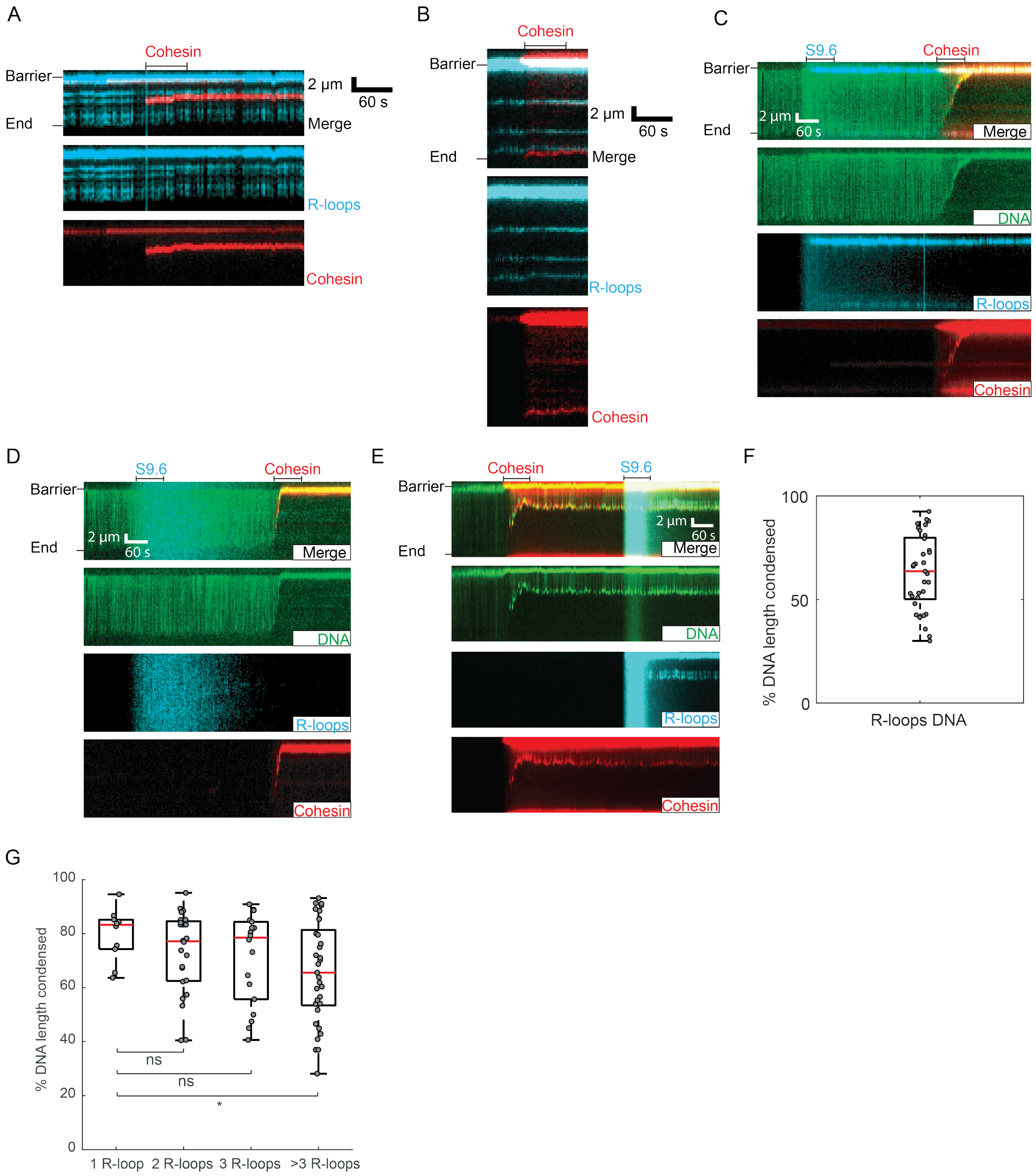


**Figure S11. Additional analysis of cohesin and R-loop collisions. Related to Figure 5.**

(A,B) Representative kymographs showing cohesin co-localized with R-loops.

(C) Representative kymograph showing that the S9.6 antibody doesn't stain any R-loops when the DNA is pre-treated with RNase H. Cohesin can also completely compact this substrate.

(D) Representative kymograph showing cohesin completely compacts non-transcribed ligated DNA.

(E) Representative kymograph showing that R-loops that are not labeled with S9.6 also slow cohesin. S9.6 was injected into the flowcell after cohesin translocation was complete.

(F) Quantification of percent DNA compacted for experiments where cohesin was injected into the flowcell prior to S9.6 injection, as shown in (E).

(G) DNA condensation as a function of the number of S9.6-stained R-loops. (N=10 for a single R-loop; N=25 for two R-loop; N=18 for triple R-loops and N=38 for >3 R-loops). Red lines: median values.

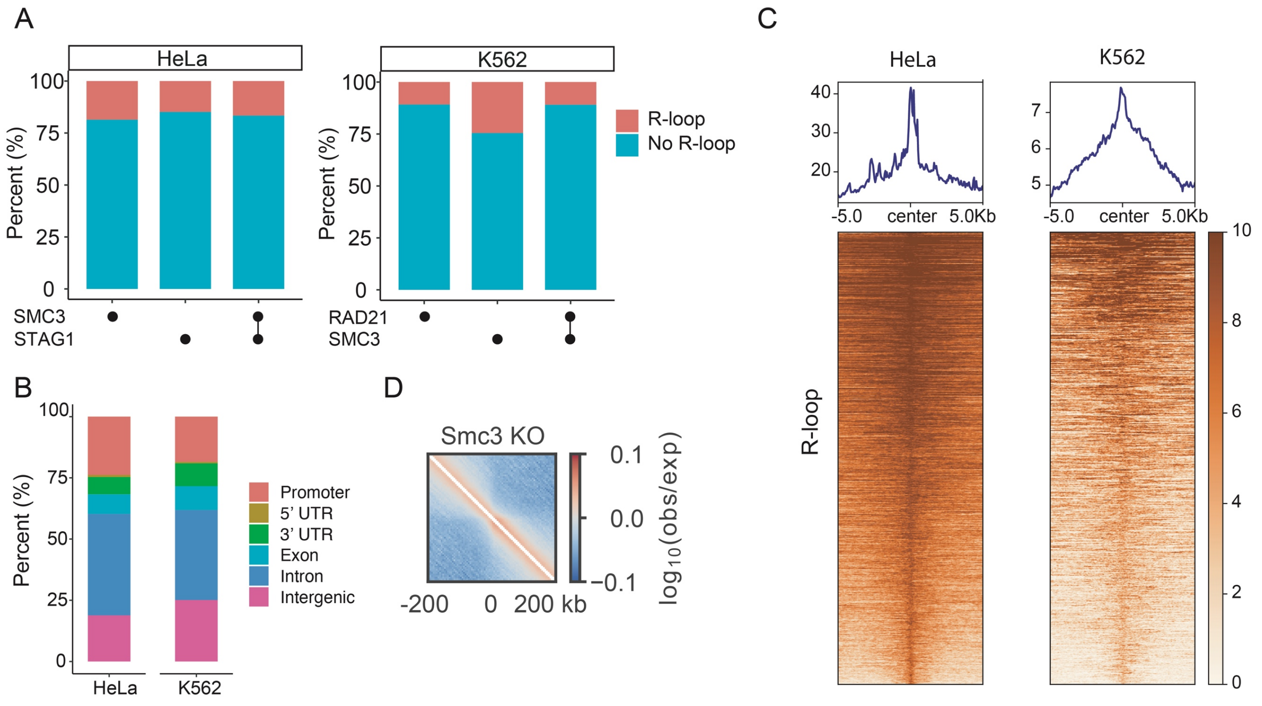


**Figure S12. R-loops co-occur with cohesin in mammalian cell lines. Related to Figure 6.**

(A) SMC3 and STAG1 positions overlap with R-loops in HeLa cells. SMC3 and RAD21 also overlap with R-loops in K562 cells.

(B) Genomic features of these overlapped peaks in HeLa and K562 cells.

(C) Read density profiles and heatmaps of R-loop reads across overlaps of SMC3, STAG1 and R-loop peaks in HeLa cells, and of RAD21, SMC3 and R-loop peaks in K562 cells.

(D) Map of average chromatin contact enrichment (observed-over-expected) in Smc3 KO MEFs centered on R-loops (n=39,680).

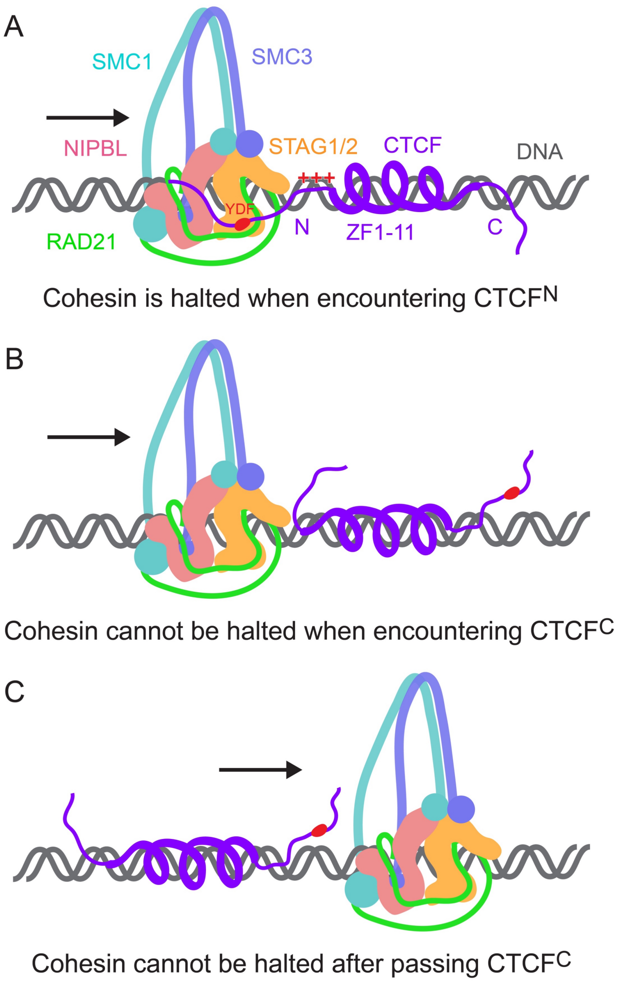


**Figure S13. Model for cohesin encounter with CTCF on DNA.**

(A) CTCF YDF motif and positively charged linker region (+++) together halt cohesin translocation on DNA.

(B-C) Cohesin can pass CTCF-bound CBSs either when encountering CTCF C-terminal side (B) or crossing over CTCF (C), owing to being incapable of interaction between the CTCF YDF motif and STAG1/2.

#### SUPPLEMENTARY MOVIES

**Movie S1.** A single-tethered DNA curtain showing CTCF-DNA homogeneously stained with SYTOX Orange bound with CTCF-Alexa488. The *cosL* end of DNA was immobilized on surface.

**Movies S2-S3.** SYTOX Orange stained U-shaped CTCF-DNA bound with CTCF-Alexa488 was extruded by cohesin-Alexa647. Restriction enzyme SfoI was loaded at the end to determine the orientation of CTCF on DNA.

**Movie S4.** $CTCF^{N}$ block cohesin translocation along DNA.

**Movie S5.** $CTCF^{C}$ accelerates cohesin translocation along DNA.

**Movie S6.** $Cas9^{Front}$ block cohesin translocation along DNA.

**Movie S7.** $Cas9^{Back}$ accelerate cohesin translocation along DNA.

**Movie S8.** R-loops block cohesion-Alexa647 translocation. The R-loops are visualized with S9.6-Alexa488 on SYTOX Orange stained DNA.

#### SUPPLEMENTARY TABLES

**Table S1. Cryo-EM data collection and model statistics. Related to Figure 3.**

| | |
|---|---|
| **Data collection and processing** | |
| Magnification | 46,296 |
| Voltage (kV) | 300 |
| Electron exposure ($e^-$/Å$^2$) | 60 |
| Defocus rage (µm) | 1.5 – 2.5 |
| Pixel size (Å) | 1.44 |
| Symmetry imposed | *C*1 |
| Initial particle images | 2,185,704 |
| Final particle images | 42,704 |
| Map resolution (Å) | 6.5 |
| FSC threshold | 0.143 |
| **Refinement** | |
| Initial model used (PDB code) | 5T0U, 5YEL, 6WGE, 6QNX |
| Model resolution (Å) | 5.8/6.2/7.1 |
| FSC threshold | 0/0.143/0.5 |
| Model Composition | |
| Non-hydrogen atoms | 34315 |
| Proteins residues | 3690 |
| Nucleotides | 216 |
| Ligands | 15 |
| B factors (Å$^2$) | |
| Protein | 240.41 |
| Nucleotide | 426.53 |
| Ligand | 178.68 |
| Bonds RMSD | |
| Length (Å) | 0.005 |
| Angles (°) | 0.929 |
| **Validation** | |
| MolProbity score | 2.09 |
| Clash score | 15.13 |
| Ramachandran plot (%) | |
| Favored | 93.87 |
| Allowed | 6.07 |
| Outliers | 0.05 |

**Table S2. DNA used in this work.**

| Name | Description | Sequence |
|---|---|---|
| dsDNA_CBS | CTCF-DNA complex purification and protein structure study | GCAAGATTGCAGTGCCCACAGAGGC<br>CAGCAGGGGGCGCTAGTGAGGTGGT<br>TTTTATATGTTTTGTTATGTATTGTTT<br>ATTTTCCCTTTAATTTTAGGATATGA<br>AAACAAGAATTTATC<br>The underlined sequence is the CTCF-binding site (CBS) |
| STAG1- W337A -F | Forward primer of PCR for STAG1 W337A | AAATACGTGGGCGCGACGCTGCACGATC GTCAGGGTG |
| STAG1- W337A -R | Reverse primer of PCR for STAG1 W337A | TCGTGCAGCGTCGCGCCCACGTATTTCA GATAGCTGTC |
| STAG1- F374A -F | Forward primer of PCR for STAG1 F374A | CTGAAATACGTGGGCGCGACGCTGCACG ATCG |
| STAG1- F374A -R | Reverse primer of PCR for STAG1 F374A | CGATCGTGCAGCGTCGCGCCCACGTATT TCAG |
| CTCF-1-F | Forward primer of PCR for CTCF | TATGGCCGGCCaATGGAAGGTGATGCAG TC |
| CTCF-727-A-NS | Reverse primer of PCR for CTCF | TTGGCGCGCCCCGGTCCATCATGCTGAG |
| CTCF_266-F | Forward primer of PCR for CTCF ZFs | TATGGCCGGCCaATGTTCCAGTGTGAGC TTTG |
| CTCF_580_NS-A | Reverse primer of PCR for CTCF ZFs | TTGGCGCGCCTGGGCCAGCACAATTATC |
| CTCF-Y226/F228A-F | Forward primer of PCR for CTCF Y226/F228A | GCAAAGATGTAGATGTGTCTGTCGCCGA TGCTGAGGAAGAACAGCAGGAGGGTC |
| CTCF-Y226/F228A-R | Reverse primer of PCR for CTCF Y226/F228A | GACCCTCCTGCTGTTCTTCCTCAGCATCG GCGACAGACACATCTACATCTTTGC |
| IF751 | Forward primer of colony PCR for recombineering check | GAA CAA ACA ATA CCC AGA TTG CG |
| IF752 | Reverse primer of colony PCR for recombineering check | GGA ATA TCT GGC GGT GCA AT |
| Lab06 | Oligo for *cosR* end annealing of CTCF$^{C}$-DNA substrate | /5Phos/GGG CGG CGA CCT/3BioTEG |

| | | |
|---|---|---|
| Lab07 | Oligo for *cosL* end annealing of CTCF$^{N}$-DNA substrate | /5Phos/AGG TCG CCG CCC/3BioTEG |
| Lab08 | Oligo for *cosL* end annealing of CTCF$^{C}$-DNA substrate | /5Phos/AGG TCG CCG CCC/3Dig_N |
| Lab09 | Oligo for *cosR* end annealing of CTCF$^{N}$-DNA substrate | /5Phos/GGG CGG CGA CCT/3Dig_N |

**Table S3. Single guide RNAs used in this work.**

| Name | RNA sequence |
|---|---|
| dCas9 sgRNA | GUG AUA AGU GGA AUG CCA UGG UUU UAG AGC UAG AAA UAG CAA GUU AAA AUA AGG CUA GUC CGU UAU CAA CUU GAA AAA GUG GCA CCG AGU CGG UGC UUU U |
| dCas12a crRNA | GUC AAA AGA CCU UUU UAA UUU CUA CUC UUG UAG AUA GGA UGA ACA GUU CUG GCU GGA GU |

**Table S4. The list of DRIP-Seq, ChIP-Seq and Hi-C data for Mouse Embryonic Fibroblasts. Related to Figure 6.**

| Mus musculus cell line | Seq type | GEO accession number | Pull-down in the indicated knockout (KO) | Citation |
|---|---|---|---|---|
| Mouse Embryonic Fibroblasts | DRIP-seq | GSE70189 | R-loop | (Sanz et al., 2016) |
| | ChIP-seq | GSM1979724 | Stag1 | (Busslinger et al., 2017) |
| | | GSM1979725 | Stag1 | |
| | | GSM1979726 | Stag1 in CTCF KO | |
| | | GSM1979727 | Stag1 in CTCF KO | |
| | | GSM1979728 | Stag1 in CTCF KO | |
| | | GSM1979732 | Stag1 in Wapl KO | |
| | | GSM1979733 | Stag1 in Wapl KO | |
| | | GSM1979734 | Stag1 in Wapl KO | |
| | | GSM1979735 | Stag1 in CTCF Wapl KO | |
| | | GSM1979736 | Stag1 in CTCF Wapl KO | |
| | | GSM1979737 | Rad21 | |
| | | GSM1979738 | Rad22 | |
| | | GSM1979739 | Rad21 in CTCF KO | |
| | | GSM1979740 | Rad21 in CTCF KO | |
| | | GSM2221806 | RAD21 in CTCF KO | |
| | | GSM1979744 | RAD21 in Wapl KO | |
| | | GSM1979745 | RAD21 in Wapl KO | |
| | | GSM1979746 | RAD21 in Wapl KO | |
| | | GSM1979747 | RAD21 in CTCF Wapl KO | |
| | | GSM1979748 | RAD21 in CTCF Wapl KO | |
| | Hi-C | GSE196621 | Not applicable | (Banigan et al., 2022) |

**Table S5. Called peaks, peak overlaps and overlap p-values of ChIP/DRIP-seq for mouse embryonic fibroblasts. Related to Figure 6.**

| **Mouse Embryonic Fibroblasts cell line** | **WT** | | | | **CTCF KO** | | | | **Wapl KO** | | | | **DKO** | | | |
|---|---|---|---|---|---|---|---|---|---|---|---|---|---|---|---|---|
| **Genotype** | Rad21 | Stag1 | Rad21 &Stag1 | R-loop | Rad21 | Stag1 | Rad21 &Stag1 | R-loop | Rad21 | Stag1 | Rad21 &Stag1 | R-loop | Rad21 | Stag1 | Rad21 &Stag1 | R-loop |
| **# Total peaks** | 20891 | 27088 | 19952 | 35894 | 37713 | 34925 | 28419 | 35894 | 26346 | 27673 | 24070 | 35894 | 12841 | 12515 | 7923 | 35894 |
| **# Peaks overlapped with R-loop (at least 1 bp in common)** | 2920 | 3952 | 2695 | NA | 8114 | 9228 | 7037 | NA | 3826 | 4064 | 3356 | NA | 2699 | 3136 | 1654 | NA |
| **P-value (Fisher's exact test, bedtools )** | < 1e-16 | < 1e-16 | < 1e-16 | NA | < 1e-16 | < 1e-16 | < 1e-16 | NA | < 1e-16 | < 1e-16 | < 1e-16 | NA | < 1e-16 | < 1e-16 | < 1e-16 | NA |
| **P-value (ChiPseeker, nShuffle=10000 )** | < 1e-4 | < 1e-4 | < 1e-4 | NA | < 1e-4 | < 1e-4 | < 1e-4 | NA | < 1e-4 | < 1e-4 | < 1e-4 | NA | < 1e-4 | < 1e-4 | < 1e-4 | NA |
| **P-value (Genomic HyperBrowser, Monte Carlo sampling=10000)** | < 1e-4 | < 1e-4 | < 1e-4 | NA | < 1e-4 | < 1e-4 | < 1e-4 | NA | < 1e-4 | < 1e-4 | < 1e-4 | NA | < 1e-4 | < 1e-4 | < 1e-4 | NA |

Note: We used the R-loop peaks called in WT cells across all KO experiments (shown in grey) because DRIP-seq data was not available for these knockouts.

**Table S6. ChIP-Seq and DRIP-Seq sample list for human cell lines. Related to Figure S12.**

<table>
<tr><th>Human cell line</th><th>Seq type</th><th>GEO accession number</th><th>Pull-down in the indicated knockout (KO)</th><th>Citation</th></tr>
<tr><td rowspan="4">HeLa</td><td rowspan="2">DRIP-seq</td><td>GSM2452072</td><td>R-loop</td><td rowspan="2">(Hamperl et al., 2017)</td></tr>
<tr><td>GSM2668157</td><td>R-loop</td></tr>
<tr><td rowspan="2">ChIP-seq</td><td>GSM3619484</td><td>SMC3</td><td rowspan="2">(Holzmann et al., 2019)</td></tr>
<tr><td>GSM3619485</td><td>EGFP-SA1</td></tr>
<tr><td rowspan="3">K562</td><td>DRIP-seq</td><td>GSM1720619</td><td>R-loop</td><td>(Sanz et al., 2016)</td></tr>
<tr><td rowspan="2">ChIP-seq</td><td>GSM935310</td><td>SMC3</td><td rowspan="2">(Pope et al., 2014)</td></tr>
<tr><td>GSM935319</td><td>RAD21</td></tr>
</table>

**Table S7. Called peaks, peak overlaps and overlap p-values of ChIP/DRIP-seq for human cell lines. Related to Figure S12.**

| **Human cell line** | **HeLa** | | | | **K562** | | | |
|---|---|---|---|---|---|---|---|---|
| **Pull-downs** | SMC3 | SA1 | SMC3 &SA1 | R-loop | RAD21 | SMC3 | RAD21 &SMC3 | R-loop |
| **# Total peaks** | 29459 | 39799 | 26099 | 89246 | 12761 | 42740 | 12609 | 50482 |
| **# Peaks overlapped with R-loop** | 5470 | 5910 | 4335 | NA | 1381 | 10467 | 1374 | NA |
| **P-value (Fisher's exact test, bedtools)** | < 1e-16 | < 1e-16 | < 1e-16 | NA | < 1e-16 | < 1e-16 | < 1e-16 | NA |
| **P-value (ChIPseeker, nShuffle=10000 )** | < 1e-4 | < 1e-4 | < 1e-4 | NA | < 1e-4 | < 1e-4 | < 1e-4 | NA |
| **P-value (Genomic HyperBrowser, Monte Carlo sampling=10000)** | < 1e-4 | < 1e-4 | < 1e-4 | NA | < 1e-4 | < 1e-4 | < 1e-4 | NA |

#### SEQUENCES OF CTCF INSERTION CASSETTES FOR RECOMBINEERING

##### Insertion cassette containing 4X CTCF binding sites

**Bold:** Homology for lambda DNA

Red: CTCF binding sites

Blue: Ampicillin resistance cassette

Restriction sites: NgoMIV cutting site is underlined.

**CGGGAATGATCCAGATTTTGCTACCACCATGACTAACGCGCTTGCGGGTAAACAACCGAAGAATGCGACACTGACGG**
**CGCTGGCAGGGCTTTCCACGGCGAAAAATAAATTACCGTATTTTGCGGAAAATGATGCCGCCAGCCTGACTGA**GCCG
GCGAGCACTCGAGTTGCATTGCAGTGCCCACAGAGGCCAGCAGGGGGCGCTAGTGAGGACCGTATTTTGCGGCCATG
GGAGGACCCAGGAGCCAGCAGAGGGCGAGGGAACAGCGCGGAACCCCTATTTGTTTATTTTTCTAAATACATTCAAA
TATGTATCCGCTCATGAGACAATAACCCTGATAAATGCTTCAATAATATTGAAAAAGGAAGAGTATGAGTATTCAAC
ATTTCCGTGTCGCCCTTATTCCCTTTTTTGCGGCATTTTGCCTTCCTGTTTTTGCTCACCCAGAAACGCTGGTGAAA
GTAAAAGATGCTGAAGATCAGTTGGGTGCACGAGTGGGTTACATCGAACTGGATCTCAACAGCGGTAAGATCCTTGA
GAGTTTTCGCCCCGAAGAACGTTTTCCAATGATGAGCACTTTTAAAGTTCTGCTATGTGGCGCGGTATTATCCCGTA
TTGACGCCGGGCAAGAGCAACTCGGTCGCCGCATACACTATTCTCAGAATGACTTGGTTGAGTACTCACCAGTCACA
GAAAAGCATCTTACGGATGGCATGACAGTAAGAGAATTATGCAGTGCTGCCATAACCATGAGTGATAACACTGCGGC
CAACTTACTTCTGACAACGATCGGAGGACCGAAGGAGCTAACCGCTTTTTTGCACAACATGGGGGATCATGTAACTC
GCCTTGATCGTTGGGAACCGGAGCTGAATGAAGCCATACCAAACGACGAGCGTGACACCACGATGCCTGTAGCAATG
GCAACAACGTTGCGCAAACTATTAACTGGCGAACTACTTACTCTAGCTTCCCGGCAACAATTAATAGACTGGATGGA
GGCGGATAAAGTTGCAGGACCACTTCTGCGCTCGGCCCTTCCGGCTGGCTGGTTTATTGCTGATAAATCTGGAGCCG
GTGAGCGTGGGTCTCGCGGTATCATTGCAGCACTGGGGCCAGATGGTAAGCCCTCCCGTATCGTAGTTATCTACACG
ACGGGGAGTCAGGCAACTATGGATGAACGAAATAGACAGATCGCTGAGATAGGTGCCTCACTGATTAAGCATTGGTA
ACTGTCAGACCAAGTTTACTCATATATACTTTAGATTGATTTAAAACTTCATTTTTAATTTAAAAGGATCTAGGTGA
AGATCCTTTTTGATAATCTCATGACCAAAATCCCTTAACGTGAGTTTTCGTTCCACTGAGCGTCAGACCCCGTAGAA
AAGATCAAAGGATCTTCTTGAGATCCTTTTTTTCTGCGCGTAATCTGCTGCTTGCAAACAAAAAAACCACCGCTACC
AGCGGTGGTTTGTTTGCCGGATCAAGAGCTACCAACTCTATTGCAGTGCCCACAGAGGCCAGCAGGGGGCGCTAGTG
AGGAAAAATAAATTACCCCATGGGAGGACCCAGGAGCCAGCAGAGGGCGAGGGAACAG**TATTGTGAGGCTTGCATAA**
**TGGCATTCAGAATGAGTGAACAACCACGGACCATAAAAATTTATAATCTGCTGGCCGGAACTAATGAATTT**
**ATTGGTGAAGGTGACGCATATATTCCGCCTCATACCGGTCTGCCTGCAAACAGTACCGAT**

**Insertion cassette containing 3X CTCF binding sites**

**Bold:** Homology for lambda DNA

Red: CTCF binding sites

Blue: Ampicillin resistance cassette

Restriction sites: NgoMIV cutting site is underlined.

**CGGGAATGATCCAGATTTTGCTACCACCATGACTAACGCGCTTGCGGGTAAACAACCGAAGAATGCGACACTGACGG**
**CGCTGGCAGGGCTTTCCACGGCGAAAAATAAATTACCGTATTTTGCGGAAAATGATGCCGCCAGCCTGACTGA**GCCG
GCGAGCACTCGAGTTGCATTGCAGTGCCCACAGAGGCCAGCAGGGGGCGCTAGTGAGGACCGTATTTTGCGGCCATG
GGAGGACCCAGGAGCCAGCAGAGGGCGAGGGAACAGCGCGGAACCCCTATTTGTTTATTTTTCTAAATACATTCAAA
TATGTATCCGCTCATGAGACAATAACCCTGATAAATGCTTCAATAATATTGAAAAAGGAAGAGTATGAGTATTCAAC
ATTTCCGTGTCGCCCTTATTCCCTTTTTTGCGGCATTTTGCCTTCCTGTTTTTGCTCACCCAGAAACGCTGGTGAAA
GTAAAAGATGCTGAAGATCAGTTGGGTGCACGAGTGGGTTACATCGAACTGGATCTCAACAGCGGTAAGATCCTTGA
GAGTTTTCGCCCCGAAGAACGTTTTCCAATGATGAGCACTTTTAAAGTTCTGCTATGTGGCGCGGTATTATCCCGTA
TTGACGCCGGGCAAGAGCAACTCGGTCGCCGCATACACTATTCTCAGAATGACTTGGTTGAGTACTCACCAGTCACA
GAAAAGCATCTTACGGATGGCATGACAGTAAGAGAATTATGCAGTGCTGCCATAACCATGAGTGATAACACTGCGGC
CAACTTACTTCTGACAACGATCGGAGGACCGAAGGAGCTAACCGCTTTTTTGCACAACATGGGGGATCATGTAACTC
GCCTTGATCGTTGGGAACCGGAGCTGAATGAAGCCATACCAAACGACGAGCGTGACACCACGATGCCTGTAGCAATG
GCAACAACGTTGCGCAAACTATTAACTGGCGAACTACTTACTCTAGCTTCCCGGCAACAATTAATAGACTGGATGGA
GGCGGATAAAGTTGCAGGACCACTTCTGCGCTCGGCCCTTCCGGCTGGCTGGTTTATTGCTGATAAATCTGGAGCCG
GTGAGCGTGGGTCTCGCGGTATCATTGCAGCACTGGGGCCAGATGGTAAGCCCTCCCGTATCGTAGTTATCTACACG
ACGGGGAGTCAGGCAACTATGGATGAACGAAATAGACAGATCGCTGAGATAGGTGCCTCACTGATTAAGCATTGGTA
ACTGTCAGACCAAGTTTACTCATATATACTTTAGATTGATTTAAAACTTCATTTTTAATTTAAAAGGATCTAGGTGA
AGATCCTTTTTGATAATCTCATGACCAAAATCCCTTAACGTGAGTTTTCGTTCCACTGAGCGTCAGACCCCGTAGAA
AAGATCAAAGGATCTTCTTGAGATCCTTTTTTTCTGCGCGTAATCTGCTGCTTGCAAACAAAAAAACCACCGCTACC
AGCGGTGGTTTGTTTGCCGGATCAAGAGCTACCAACTCTATTGCAGTGCCCACAGAGGCCAGCAGGGGGCGCTAGTG
AGGAAAAATAAATTACC**TATTGTGAGGCTTGCATAATGGCATTCAGAATGAGTGAACAACCACGGACCATAAAAATT**
**TATAATCTGCTGGCCGGAACTAATGAATTTATTGGTGAAGGTGACGCATATATTCCGCCTCATACCGGTCTGCCTGC**
**AAACAGTACCGAT**

**Insertion cassette containing 2X CTCF binding sites**

**Bold:** Homology for lambda DNA

Red: CTCF binding sites

Blue: Ampicillin resistance cassette

Restriction sites: NgoMIV cutting site is underlined.

**CGGGAATGATCCAGATTTTGCTACCACCATGACTAACGCGCTTGCGGGTAAACAACCGAAGAATGCGACACTGACGG**
**CGCTGGCAGGGCTTTCCACGGCGAAAAATAAATTACCGTATTTTGCGGAAAATGATGCCGCCAGCCTGACTGA**GCCG
GCGAGCACTCGAGTTGCATTGCAGTGCCCACAGAGGCCAGCAGGGGGCGCTAGTGAGGACCGTATTTTGCGGCGCGG
AACCCCTATTTGTTTATTTTTCTAAATACATTCAAATATGTATCCGCTCATGAGACAATAACCCTGATAAATGCTTC
AATAATATTGAAAAAGGAAGAGTATGAGTATTCAACATTTCCGTGTCGCCCTTATTCCCTTTTTTGCGGCATTTTGC
CTTCCTGTTTTTGCTCACCCAGAAACGCTGGTGAAAGTAAAAGATGCTGAAGATCAGTTGGGTGCACGAGTGGGTTA
CATCGAACTGGATCTCAACAGCGGTAAGATCCTTGAGAGTTTTCGCCCCGAAGAACGTTTTCCAATGATGAGCACTT
TTAAAGTTCTGCTATGTGGCGCGGTATTATCCCGTATTGACGCCGGGCAAGAGCAACTCGGTCGCCGCATACACTAT
TCTCAGAATGACTTGGTTGAGTACTCACCAGTCACAGAAAAGCATCTTACGGATGGCATGACAGTAAGAGAATTATG
CAGTGCTGCCATAACCATGAGTGATAACACTGCGGCCAACTTACTTCTGACAACGATCGGAGGACCGAAGGAGCTAA
CCGCTTTTTTGCACAACATGGGGGATCATGTAACTCGCCTTGATCGTTGGGAACCGGAGCTGAATGAAGCCATACCA
AACGACGAGCGTGACACCACGATGCCTGTAGCAATGGCAACAACGTTGCGCAAACTATTAACTGGCGAACTACTTAC
TCTAGCTTCCCGGCAACAATTAATAGACTGGATGGAGGCGGATAAAGTTGCAGGACCACTTCTGCGCTCGGCCCTTC
CGGCTGGCTGGTTTATTGCTGATAAATCTGGAGCCGGTGAGCGTGGGTCTCGCGGTATCATTGCAGCACTGGGGCCA
GATGGTAAGCCCTCCCGTATCGTAGTTATCTACACGACGGGGAGTCAGGCAACTATGGATGAACGAAATAGACAGAT
CGCTGAGATAGGTGCCTCACTGATTAAGCATTGGTAACTGTCAGACCAAGTTTACTCATATATACTTTAGATTGATT
TAAAACTTCATTTTTAATTTAAAAGGATCTAGGTGAAGATCCTTTTTGATAATCTCATGACCAAAATCCCTTAACGT
GAGTTTTCGTTCCACTGAGCGTCAGACCCCGTAGAAAAGATCAAAGGATCTTCTTGAGATCCTTTTTTTCTGCGCGT
AATCTGCTGCTTGCAAACAAAAAAACCACCGCTACCAGCGGTGGTTTGTTTGCCGGATCAAGAGCTACCAACTCTAT
TGCAGTGCCCACAGAGGCCAGCAGGGGGCGCTAGTGAGGAAAAATAAATTACC**TATTGTGAGGCTTGCATAATGGCA**
**TTCAGAATGAGTGAACAACCACGGACCATAAAAATTTATAATCTGCTGGCCGGAACTAATGAATTTATTGGTGAAGG**
**TGACGCATATATTCCGCCTCATACCGGTCTGCCTGCAAACAGTACCGAT**

**Insertion cassette containing 1X CTCF binding sites**

**Bold:** Homology for lambda DNA

Red: CTCF binding sites

Blue: Ampicillin resistance cassette

Restriction sites: NgoMIV cutting site is underlined.

**CGGGAATGATCCAGATTTTGCTACCACCATGACTAACGCGCTTGCGGGTAAACAACCGAAGAATGCGACACTGACGG**
**CGCTGGCAGGGCTTTCCACGGCGAAAAATAAATTACCGTATTTTGCGGAAAATGATGCCGCCAGCCTGACTGA**GCCG
GCGAGCACTCGAGTTGCTAATTGCAGTGCCCACAGAGGCCAGCAGGGGGCGCTAGTGAGGTTGGACGCGGAACCCCT
ATTTGTTTATTTTTCTAAATACATTCAAATATGTATCCGCTCATGAGACAATAACCCTGATAAATGCTTCAATAATA
TTGAAAAAGGAAGAGTATGAGTATTCAACATTTCCGTGTCGCCCTTATTCCCTTTTTTGCGGCATTTTGCCTTCCTG
TTTTTGCTCACCCAGAAACGCTGGTGAAAGTAAAAGATGCTGAAGATCAGTTGGGTGCACGAGTGGGTTACATCGAA
CTGGATCTCAACAGCGGTAAGATCCTTGAGAGTTTTCGCCCCGAAGAACGTTTTCCAATGATGAGCACTTTTAAAGT
TCTGCTATGTGGCGCGGTATTATCCCGTATTGACGCCGGGCAAGAGCAACTCGGTCGCCGCATACACTATTCTCAGA
ATGACTTGGTTGAGTACTCACCAGTCACAGAAAAGCATCTTACGGATGGCATGACAGTAAGAGAATTATGCAGTGCT
GCCATAACCATGAGTGATAACACTGCGGCCAACTTACTTCTGACAACGATCGGAGGACCGAAGGAGCTAACCGCTTT
TTTGCACAACATGGGGGATCATGTAACTCGCCTTGATCGTTGGGAACCGGAGCTGAATGAAGCCATACCAAACGACG
AGCGTGACACCACGATGCCTGTAGCAATGGCAACAACGTTGCGCAAACTATTAACTGGCGAACTACTTACTCTAGCT
TCCCGGCAACAATTAATAGACTGGATGGAGGCGGATAAAGTTGCAGGACCACTTCTGCGCTCGGCCCTTCCGGCTGG
CTGGTTTATTGCTGATAAATCTGGAGCCGGTGAGCGTGGGTCTCGCGGTATCATTGCAGCACTGGGGCCAGATGGTA
AGCCCTCCCGTATCGTAGTTATCTACACGACGGGGAGTCAGGCAACTATGGATGAACGAAATAGACAGATCGCTGAG
ATAGGTGCCTCACTGATTAAGCATTGGTAACTGTCAGACCAAGTTTACTCATATATACTTTAGATTGATTTAAAACT
TCATTTTTAATTTAAAAGGATCTAGGTGAAGATCCTTTTTGATAATCTCATGACCAAAATCCCTTAACGTGAGTTTT
CGTTCCACTGAGCGTCAGACCCCGTAGAAAAGATCAAAGGATCTTCTTGAGATCCTTTTTTTCTGCGCGTAATCTGC
TGCTTGCAAACAAAAAAACCACCGCTACCAGCGGTGGTTTGTTTGCCGGATCAAGAGCTACCAACTCT**TATTGTGAG**
**GCTTGCATAATGGCATTCAGAATGAGTGAACAACCACGGACCATAAAAATTTATAATCTGCTGGCCGGAACTAATGA**
**ATTTATTGGTGAAGGTGACGCATATATTCCGCCTCATACCGGTCTGCCTGCAAACAGTACCGAT**

#### METHODS AND MATERIALS

##### Protein expression and purification

Cohesin-NIPBL complex

The human cohesin-NIPBL complex was expressed and purified as described previously (Kim et al., 2019; Shi et al., 2020). Briefly, SMC1, SMC3, RAD21 (R172/D279/R450A) with a C-terminal MBP tag, and $NIPBL^C$ (residues 1163-2804) with an N-terminal His-MBP tag were co-expressed in High Five cells (Thermo Fisher). SMC1, SMC3, RAD21 and $NIPBL^C$ (cohesin- $NIPBL^C$ without STAG1) were co-purified by Amylose Resin (NEB), a HiTrap Heparin HP column (GE Healthcare), a Resource Q column and a Superose 6 Increase 10/300 GL column (GE Healthcare). STAG1 with a C-terminal TwinStrepII-His dual tag or with a SNAPf-TwinStrepII-His tag was expressed separately. STAG1 was purified by $Ni^{2+}$-NTA agarose (Qiagen) followed by a Resource Q column (GE Healthcare). The C-terminal dual tag in STAG1 was removed by home-made 3C protease. To form complete cohesin-$NIPBL^C$ complex, STAG1 protein was added before Heparin column or Superose 6 Increase 10/300 GL column. Cohesin mutants, cohesin-WFA and cohesin-EQ were purified as wildtype proteins. Purified complexes were concentrated, flash frozen in liquid nitrogen and stored at -80 °C.

CTCF

Full length CTCF was cloned into a modified pFastbac Dual vector (Thermo Fisher) with a C-terminal MBP and Flag dual tag and expressed in High Five cells. Cells were resuspended in Lysis Buffer TN500 (20 mM Tris pH 7.5, 500 mM NaCl, 5% glycerol, 5 mM 2-mercaptoethanol) plus 25 µM $ZnSO_4$, 1 × Pierce™ EDTA-free Protease Inhibitor Tablets (Thermo Fisher), and 1 mM phenylmethylsulfonyl fluoride (PMSF), and lysed by a high-pressure homogenizer (Microfluidics LM-20). After the addition of 0.3% polyethylenimine, cell lysate was cleared by centrifugation at

40,000 g for 1 h at 4˚C. The supernatant was mixed with Amylose Resin for 1.5 h in the cold room. The resin was washed with Lysis Buffer TN500 plus 500 mM NaCl and 25 μM $ZnSO_4$, followed by the Buffer TN150 (20 mM Tris pH 8.0, 150 mM NaCl, 5% glycerol, 5 mM 2-mercaptoethanol) plus 25 μM $ZnSO_4$, and eluted by the Buffer TN150 plus 40 mM maltose and 25 μM $ZnSO_4$. The protein was further purified by a Resource Q column and finally a Superose 6 Increase 10/300 GL column.

To reconstitute the CTCF-DNA complex, purified CTCF and dsDNA (GCAAGATTGCAGTGCCCACAGAGGCCAGCA GGGGGCGCTAGTGAGGTGGTTTTATATGTTTTGTTATGTATTGTTTATTTTCCCTTTA ATTTTAGGATATGAAAACAAGAATTTATC; underlined sequence is CTCF-binding site, CBS) were incubated at equal molar ratio on ice for 1 h, and then loaded onto a Superose 6 Increase 10/300 GL. Purified samples were concentrated, flash-frozen in liquid nitrogen and stored at -80˚C. CTCF mutants, CTCF-YFA, CTCF-ZF (266-580) were purified as wildtype proteins.

Nuclease dead (dCas9)

The fusion construct of nuclease-dead *S. pyogenes* Cas9 (dCas9) with D10A/H840A mutations contained an N-terminal hexahistidine-maltose binding protein ($His_6$-MBP) tag and a peptide sequence containing a tobacco etch virus (TEV) protease cleavage site (Jinek et al., 2012), followed by triple Flag epitope tag. Protein was expressed in BL21 star (DE3) cells (Thermo Fisher Scientific) and purified as described previously (Jones et al., 2021). Protein expression was induced with 1 mM IPTG for 18 h at 18 °C after cells were grown to $OD_{600}$ of 0.6 at 30 °C in 1L LB medium with 50 mg $mL^{-1}$ kanamycin. Cells were collected by centrifugation and lysed by sonication at 4 °C in lysis buffer (20 mM Tris-Cl, pH 8.0, 250 mM NaCl, 5 mM imidazole, 5 μM PMSF, 6 units $mL^{-1}$ DNAse I). The lysate was clarified by ultracentrifugation at 35,000 relative centrifugal force (RCF) and then passed over a nickel affinity column (HisTrap FF 5 mL, GE Healthcare) and eluted with elution buffer (20 mM Tris-Cl, pH 8.0, 250 mM NaCl, 250 mM imidazole). The $His_6$-MBP was cleaved overnight in dialysis buffer (20 mM HEPES-KOH, pH 7.5, 150 mM KCl, 10% glycerol, 1 mM DTT, 1 mM EDTA) in the presence of home made TEV protease (0.5 mg per 50 mg of protein). The dialyzed protein was resolved on a HiTrap SP FF 5 mL column (GE Healthcare) with a linear gradient between buffer A (20 mM HEPES-KOH, pH 7.5, 100 mM KCl) and buffer B (20 mM HEPES-KOH, pH 7.5, 1 M KCl). Protein-containing fractions were concentrated via dialysis (10 kDa Slide-A-Lyzer, Thermo Fisher Scientific) and then passed over a Superdex 200 Increase 10/300 column (GE Healthcare) pre-equilibrated into storage buffer (20 mM HEPES-KOH, pH 7.5, 500 mM KCl). The protein was snap frozen in liquid nitrogen and stored in 10 µL aliquots at -80 °C.

<u>Nuclease-dead Cas12a (Cas12a)</u>

The nuclease-dead *Acidaminococcus sp.* Cas12a (dCas12a) containing D908A point mutant was expressed and purified using a previously established protocol with minor modifications (Strohkendl et al., 2018). The construct of dCas12a contained a $His_6$-Twin-Strep-SUMO N-terminal fusion and a triple Flag epitope C-terminal fusion was transformed and expressed in BL21 star (DE3) cells. A 20 mL culture of Terrific Broth (TB) with 50 mg $mL^{-1}$ carbenicillin was inoculated with a single colony and grown overnight at 37 °C with shaking. 1 L of TB with 50 mg $mL^{-1}$ carbenicillin was inoculated with 10 mL of the starter culture and then grown to an $OD_{600}$ of 0.6 at 37 °C. Protein expression was induced with 0.5 mM IPTG for 24 h at 18 °C. Cells were collected by centrifugation and lysed by sonication at 4 °C in lysis buffer (20 mM Na-HEPES, pH 8.0, 1 M NaCl, 1 mM EDTA, 5% glycerol, 0.1% Tween-20, 1 mM PMSF, 2000 U DNase (GoldBio), 1× HALT protease inhibitor (Thermo Fisher Scientific)). The supernatant clarified by ultracentrifugation at 35,000 RCF was loaded to a hand-packed StrepTactin Superflow gravity column (IBA Lifesciences) and then eluted (20 mM Na-HEPES, 1 M NaCl, 5 mM desthiobiotin, 5 mM MgCl2, 5% glycerol). The eluate was concentrated using a 30-kDa MWCO spin concentrator (Millipore), and 3 μM SUMO protease was added and then incubated overnight on a rotator at 4 °C. The protein was then fractionated over a HiLoad 16/600 Superdex 200 Column (GE Healthcare) pre-equilibrated with storage buffer (20 mM Na-HEPES, 150 mM KCl, 5 mM MgCl2, 2 mM DTT buffer). The protein was finally snap frozen in liquid nitrogen and stored in 10 μL aliquots at -80 °C.

**Preparation of DNA substrates with CTCF binding sites**

The DNA substrate is derived from bacteriophage λ. We recombineered λ phage lysogens in *E. coli* using the Red system as described previously with the following modification (Kim et al., 2017). DNA cassettes withh up to four CTCF binding sites were ordered from Integrated DNA Technologies (IDT). A 5 mL LB culture of lysogen cells (IF189) transformed with pKD78 plasmid (pIF284) containing Red recombinase system were grown overnight at 30°C in the presence of 10 μg $\mu L^{-1}$ chloramphenicol. 350 μL of cells were used to inoculate a fresh 35 mL culture of LB containing the same concentration of antibiotic. When the cells reached an $OD_{600}$ ~ 0.5, the Red recombinase system was induced by adding 2% L-arabinose (GoldBio) and incubated for an additional 1 hour at 30°C. Cells were harvested at 4,500 RCF for 7 min, washed three times in ice-cold Milli-Q $H_2O$, and finally resuspended in 200 μL of $H_2O$. The cells were kept on ice and used immediately for the recombineering reaction.

For recombineering, the resuspended lysogen cells were mixed with 100 ng of the insertion cassette and were electroporated at 12.5 kV $cm^{-1}$ in 0.2 cm cuvette using a MicroPulser

Electroporator (Bio-Rad 165-210). Cells were immediately resuspended in 1 mL of SOC and then transferred to a culture tube containing 2 mL LB broth. After at least 6 hours outgrowth at 30 °C, 300 μL of the culture was plated onto LB agar plates containing 30 μg $mL^{-1}$ carbenicillin. Successful incorporation of recombinant DNA was checked via colony PCR with oligos IF751 (GAA CAA ACA ATA CCC AGA TTG CG) and oligo IF752 (GGA ATA TCT GGC GGT GCA AT) and were further confirmed by DNA sequencing.

To purify these DNA substrates from recombinant phages, we induced phage production via heat shock. A single lysogenic colony was grown in 50 mL of LB broth with 50 μg $mL^{-1}$ carbenicillin overnight at 30°C. 5 mL of this starter culture was used to inoculate 500 mL of LB on the following day. When the cells reached an $OD_{600}$ ~ 0.6, they were incubated in a 70 °C water bath to rapidly increase the temperature to 42 °C. The culture was placed at 45 °C in a shaking incubator for 15 minutes and then transferred to a 37 °C incubator for 2 hours. To liberate the phage particles, cells were harvested by centrifugation at 3,000 RCF for 30 minutes and lysed via resuspension in 10 mL of SM buffer (50 mM Tris-HCl pH 7.5, 100 mM NaCl, 8 mM $MgSO_4$) with 2% chloroform, and rotated at 37 °C for 30 min. A subsequent 1 hour incubation with 50 ng $\mu L^{-1}$ DNase I (Sigma-Aldrich D2821) and 30 ng $\mu L^{-1}$ RNaseA (Sigma-Aldrich R6513) degraded the bacterial genomic DNA and RNA. The clarified lysate, containing soluble phage capsids, was obtained by centrifugation for 15 minutes at 6,000 RCF at 4 °C, and further diluted with 40 mL of SM buffer. Phage capsids were precipitated by incubating for 1 hour with 10 mL ice-cold buffer L2 (30% PEG 6000, 3 M NaCl) and then harvested by centrifugation at 10,000 RCF for 10 minutes at 4 °C. The phage pellet was washed with 1 mL of buffer L3 (100 mM Tris-HCl pH 7.5, 100 mM NaCl, 25 mM EDTA) and then resuspended with 3 mL of buffer L3, followed by an equal volume of buffer L4 (4% SDS). The phage capsid proteins were further digested by incubation with 100 ng $\mu L^{-1}$ of proteinase K (NEB #P8012S) for 1 hour at 55 °C. SDS was precipitated with 3 mL buffer L5 (3 M potassium acetate pH 7.5), and the cloudy solution was clarified by centrifugation at 15,000 RCF for 30 minutes at 4 °C. The soluble phage DNA was passed over a pre-equilibrated Qiagen tip-500 column (Qiagen #10262), washed with buffer QC (1.0 M NaCl, 50 mM MOPS pH 7.0, 15% isopropanol) and eluted with 15 mL buffer QF (1.25 M NaCl, 50 mM Tris pH 8.5, 15% isopropanol). Finally, DNA was precipitated with the addition of 10.5 mL of 100% isopropanol, rinsed in 70% ethanol twice and dissolved in 500 µL TE buffer (10 mM Tris-HCl pH 8.0, 1 mM EDTA). DNA was stored at 4 ˚C.

Biotinylation of the DNA substates for single-molecule microscopy

To prepare DNA substates for microscopy, 125 µg of λ-phage CTCF-DNA was mixed with two oligos (2 µM oligo Lab07 and 2 µM oligo Lab09 for $CTCF^{N}$-DNA, or 2 µM / oligo Lab06 and 2 µM oligo Lab08 for $CTCF^{C}$-DNA) in 1× T4 DNA ligase reaction buffer (NEB B0202S) and heated to 70°C for 15 min followed by gradual cooling to 15°C for 2 hours. One oligo will be annealed with the overhand located at the left cohesive end (*cosL*) of DNA, and the other oligo will be annealed with the overhand at right cohesive end (*cosR*). After the oligomer hybridization, 2 µL of T4 DNA ligase (NEB M0202S) was added to the mixture and incubated overnight at room temperature to seal nicks on DNA. The ligase was inactivated with 2 M NaCl, and the reaction was injected to an S-1000 gel filtration column (GE) to remove excess oligonucleotides and proteins.

**Single-Molecule Fluorescence Microscopy**

Flowcell preparation

Flowcells used for single-molecule DNA experiments were prepared as previously described (Zhang et al., 2020). Briefly, a 4-mm-wide, 100-μm-high flow channel was constructed between a glass coverslip (VWR 48393 059) and a custom-made quartz microscope slide patterned with 1-2 μm Chromium barriers using two-sided tape (3M 665). The surface was passivated with a fluid biotinylated lipid bilayer. All experiments were conducted at 37 °C under indicated flow rate. Single-molecule fluorescent images were collected with a customized prism TIRF microscopy-based inverted Nikon Ti-E microscope system. The sample was illuminated with a 488 nm laser (Coherent Sapphire) and a 637 nm laser (Coherent OBIS) through a quartz prim (Tower Optical Co.). For imaging SYTOX Orange-stained DNA and Alexa488-labled CTCF, the 488 nm laser power was adjusted to deliver low power (4 mW) at the front face of the prism using a neutral density filter set (Thorlabs). For imaging Alexa647-labeled cohesin, the 637 nm laser power was adjusted to 10 mW. Multi-color imaging was recorded using electron-multiplying charge-coupled device (EMCCD) cameras (Andor iXon DU897). DNA was visualized by a continuous flow (0.12 mL $min^{-1}$) in the imaging buffer (40 mM Tris-HCl pH 8.0, 2 mM MgCl2, 0.2 mg $mL^{-1}$ BSA, 50 mM NaCl, 1 mM DTT, and 1 mM ATP, 100 nM SYTOX Orange) supplemented with an oxygen scavenging system (3% D-glucose (w/v), 1 mM Trolox, 1500 units catalase, 250 units glucose oxidase; all from Sigma-Aldrich). Unless indicated, 1 mM ATP was added in the imaging buffer. NIS-Elements software (Nikon) was used to collect the images at 2-3 s frame rate with 150 ms exposure time. All images were exported as uncompressed TIFF stacks for further analysis in FIJI (NIH) and MATLAB (The MathWorks).

Preparation of U-shaped CTCF-DNA substrates

The strategy of making U-shaped double-tethered DNA was described previously (Kim et al., 2019). Briefly, Two biotinylated oligos (2 μM oligo Lab06 and 2 μM oligo Lab07) were annealed and ligated to CTCF-DNA as mentioned above. The biotin-BSA stock solution (10 mg $mL^{-1}$, Sigma-Aldrich A8549) was prepared in the T50 buffer (10 mM Tris, 50 mM NaCl, pH 8.0) and stored at 4℃. The solution was diluted 10 times with the T50 buffer right before use and was injected into the flowcell. After 10 minutes BSA coating, 300 μL streptavidin solution (0.1 mg $mL^{-1}$, Invitrogen 434301) was loaded into the flowcell and incubated for 10 minutes at room temperature. For further passivation of the surface to prevent nonspecific adsorption of nucleic acids and proteins, 30 μl of the liposome stock solution (Avanti Polar Lipids; 98 mol% DOPC, 2 mol% DOPE-mPEG2K) was diluted in 800 μl of the lipid buffer (10 mM Tris pH 8.0, 100 mM NaCl) and loaded into the flowcell. After 20 minutes incubation, 20 μl of CTCF-DNA stock was diluted in 1 mL T50 buffer and incubated for 5 min in the chamber. The flowcell was then immediately used for the single-molecule imaging on the TIRF microscope.

**Preparation of R-loops DNA substrates**

The template DNA used for R-loop DNA substrates, pFC53-Airn plasmid (3991 bp), containing the mouse Airn sequence with downstream of a T3 promoter, was a generous gift from the Chedin lab (Stolz et al., 2019). The process of making long R-loop DNA was based on the previous protocol with minor modifications (Pan et al., 2020). Briefly, 600 μg template DNA plasmid was linearized with 4 μL restriction enzyme BsaI (NEB), which was then inactivated by heat. The linear DNA was ligated using Quick Ligation Kit (NEB) for overnight incubation at room temperature to form long ligated DNA. Biotinylated nucleotide was synthesized to one end of long ligated DNA. For the reaction, 25 μM dATP (NEB), 25 μM dTTP (NEB), 25 μM dGTP (NEB), 5 μM dCTP (NEB), 20 μM bio-dCTP (Jena Bioscience, NU-809-BIO-16-S) and 2 units Klenow Fragment 3′→5′ exo- (NEB M0212S) were added into the ligated DNA product in the presence of 1X T4 DNA ligase buffer (NEB B0202S). After 30 minutes incubation at 25°C, the DNA was purified using phenol/chloroform extraction and ethanol precipitation. The purified biotinylated long ligated DNA was stored at -20 °C. The long R-loops DNA was generated by *in vitro* transcription, which was carried out at 37 °C for 1 hour in 1X Transcription Optimized Buffer (Promega, P4024) with 120 μg biotinylated long ligated DNA, 4 μL T3 RNA polymerase (Promega, P4024), 20 mM DTT, 0.05% Tween-20, and 500 μM rNTP. Transcription was terminated by heat inactivation at 65 °C for 10 minutes. Before R-loops DNA was injected into flowcell, 1 μL RNase A (NEB T3018L) was added to degrade free RNA at 37 °C for 30 minutes incubation. For control DNA without R-loops, R-loops DNA was treated by adding 2 μL RNase H (NEB M0297S) at 37 °C for 30 minutes incubation. This protocol produces short biotinylated fragments that decoare the flowcell surface, bind cohesin, and reduce the signal to background for longer R-loop concatamers. To reduce the background signal from short biotinylated DNA fragments, 1 μm diameter streptavidin-coated beads with (NEB S1420S) were injected into the flowcell prior to R-loop DNA injection. Longer DNA molecules that bind these beads are easily distinguishable from short DNA fragments. By keeping these long DNAs on the surface, we reduce the background signal from surface-bound short DNAs. Beads that had two or more DNA molecules were not included in our analysis. We only focus on the single R-loop DNA molecule. Monoclonal antibody S9.6 (gift from Paull Lab) coupled with Goat anti-mouse Alexa Fluor 488 (Thermo Fisher, A28175) was used to detect R-loop.

**Fluorescent imaging of proteins on DNA**

Imaging cohesin and CTCF on DNA

For imaging Alexa647-labeled wild type cohesin, 1 μM of the protein complex was diluted to 7 nM in a total volume of 150 μL imaging buffer. For the fluorescent labeling of cohesin mutants, cohesin-WFA and cohesin-EQ, 7 nM of protein were pre-incubated with 10 nM anti-$His_6$ antibody (Takara, 631212) and 12 nM anti-mouse quantum dots ($QD_{705}$) (Thermo Fisher, Q11062MP) on ice for 10 minutes. The mixture was then diluted to a total volume of 150 μL imaging buffer. For the fluorescent labeling of CTCF, CTCF-ZF, CTCF-YFA, 2 nM of protein were conjugated with 4 nM monoclonal anti-Flag antibody (Sigma Aldrich, F3164) and 6 nM Goat anti-mouse Alexa Fluor 488 on ice for 10 minutes. The mixture was then diluted to a total volume of 150 μL imaging buffer. Immediately after the conjugation or dilution, the fluorescently labeled proteins were injected into the flowcell at a 0.12 mL $min^{-1}$ flow rate. 100 μL of 5 ng/mL Heparin was loaded into the flowcell prior to CTCF to remove non-specific CTCF binding.

dCas9-RNP and dCas12a-RNP preparation and labeling

dCas9 and dCas12 ribonucleoprotein (RNP) complexes were reconstituted by incubating a 1:2 molar ratio of apoprotein and RNA (see **Table S3** for sgRNA sequences) in incubation buffer (50 mM Tris-HCl pH 8.0, 5 mM $MgCl_2$, 2 mM DTT) followed by incubation at 25 °C for 25 minutes. dCas9 and dCas12a were labeled by anti-Flag coupled with Alexa Flour 488, and then diluted to 5 nM-10 nM in imaging buffer before injected into the flowcell.

**Fluorescent image analysis**

The DNA compaction percentage and rate were analyzed in FIJI. The Mann-Whitney t-test was used to determine the significant difference between experimental conditions. Error bars on the binding distribution histogram were calculated in MATLAB using bootstrap analysis with replacement. The significance threshold was set at 0.05 in all tests.

The number of CTCF on CBS on CTCF-DNA was calculated by dividing the initial fluorescence intensity of each CTCF patch on CBS, which was averaged from first 5 frames within a 6 x 6 pixel region of interest (ROI), by the average initial intensity of CTCF patches on non-specific sites. To determine CTCF lifetime on DNA, we measured the total amount of time that each CTCF punctum spent on its DNA-binding site. Survival data were fit to single-exponential decays via a custom Matlab script (Myler et al., 2016). The boxplots plotted by Matlab denote the first and third quartiles of the data. The whiskers each cover 25% of the data values.

**Cryo-electron microscopy (Cryo-EM)**

Sample preparation and data acquisition

For cryo-EM sample preparation, purified cohesin-NIPBL$^{C}$ complex and CTCF-dsDNA complex were incubated in 20 mM HEPES pH 7.5, 50 mM KCl, 5% glycerol, 1 mM DTT, 2 mM $MgCl_2$, 0.01% IGEPAL® CA-630 for 30 min on ice. Then 1 mM ADP, 1 mM $BeSO_4$ and 8 mM NaF were added, and the sample was incubated at 30°C for 25 min. For crosslinked sample, a further 1 mM bis(sulfosuccinimidyl)suberate ($BS^3$) was added to the sample and incubated for 30 min on ice. Both samples were applied to a 20-40% sucrose gradient in 20 mM HEPES pH 7.5, 100 mM KCl, 0.5 mM TCEP, 2 mM $MgCl_2$, 0.01% IGEPAL® CA-630, and ultracentrifugation was carried out at 36,000 rpm for 16 h at 4°C. Samples were analyzed by 6% SDS-PAGE. The fractions containing cohesin-NIPBL-CTCF-DNA complex were pooled and concentrated. Then the sample buffer was exchanged to 20 mM HEPES pH 7.5, 100 mM KCl, 0.5 mM TCEP, 2 mM $MgCl_2$, 2 mM fluorinated Fos-Choline-8.

The freshly purified, crosslinked cohesin-NIPBL$^{C}$-CTCF-DNA complex ($OD_{280}$ = 7.5-10, $OD_{260}/OD_{280}$ = 1.25) was applied to glow-discharged Quantifoil R1.2/1.3 300-mesh gold holey carbon grids (Ted Pella). Grids were blotted for 2.5 s under 100% humidity at 4 °C before being plunged into liquid ethane using a Vitrobot Mark IV (FEI). Micrographs were acquired on a Titan Krios microscope (FEI) operated at 300 kV with a K3 Summit direct electron detector (Gatan), using a slit width of 20 eV on a GIF-Quantum energy filter. SerialEM software (Mastronarde, 2005) was used for automated data collection following standard procedures. A calibrated magnification of 46,296 X was used for imaging, yielding a pixel size of 1.08 Å on images. Before image processing, the images were binned further to reduce the particles image size, resulting the final pixel size of 1.44 Å. The defocus range was set from -1.5 μm to -2.5 μm. Each micrograph was dose-fractionated to 36 frames, with a total dose of about 60 $e^{-}/Å^{2}$.

Cryo-EM image processing

The detailed image processing statistics are summarized in Figs. S5 and S6 and Table S1. Motion correction was performed using the MotionCorr2 program (Zheng et al., 2017), and the CTF parameters of the micrographs were estimated using the Gctf program (Zhang, 2016). Most of steps of image processing were performed using RELION-3 (Scheres, 2012; Zivanov et al., 2018). Totally 9,121 movie frames were collected. Initially, particles picking was performed by crYOLO (Wagner et al., 2019). Class averages representing projections in different orientations selected from the initial 2D classification were used as templates for the reference based automatic particle picking. Extracted particles were binned 2 times and subjected to 2D classification. Particles from the classes with fine structural features were selected for 3D classification and reported cohesin-NIPBL-DNA structure was used as an initial model. Particles from one 3D classes showing good structural features were selected for second round classification with fine angular sampling and local search. The particles from the three good classes were chosen and subject to 3D refinement, CTF refinement and particle polishing, generating three maps with overall resolution of 3.7 Å, 7.0 Å and 6.5 Å, respectively.

To improve the resolution for CTCF and bound DNA, we performed focused 3D refinement with different sizes of soft masks. All resolutions were estimated by applying a soft mask around the protein density using the gold-standard Fourier shell correlation (FSC) = 0.143 criterion (Scheres and Chen, 2012). The local resolution was calculated with RELION-3.

Cryo-EM Model building and refinement

Models of STAG1-RAD21-CTCF and CTCF ZF1 for docking into cryo-EM maps were generated by SWISS-MODEL server (Waterhouse et al., 2018) using the structures of STAG2-RAD21-CTCF and CTCF ZFs-DNA complexes (PDB codes 6QNX and 5YEL) as templates (Hashimoto et al., 2017; Li et al., 2020; Yin et al., 2017). These models and reported structures of cohesin-NIPBL$^{C}$-DNA (PDB code 6WGE) and CTCF ZFs-DNA (PDB codes 5T0U and 5YEL) complexes (Hashimoto et al., 2017; Shi et al., 2020; Yin et al., 2017) were docked onto cryo-EM maps in UCSF Chimera (Pettersen et al., 2004) and refined by real-space refinement in Phenix with rigid body and secondary structure restraints (Adams et al., 2010; Afonine et al., 2018a). Then model was iteratively manually built in coot (Emsley et al., 2010) and refined in Phenix. Model validation was performed in Molprobity (Afonine et al., 2018b; Williams et al., 2018). Structural images were generated in UCSF ChimeraX (Goddard et al., 2018) and PyMOL (Schrodinger).

**ChIP-seq and DRIP-seq analysis**

Cohesin subunits ChIP-Seq dataset and R-loop DRIP-Seq dataset for mouse embryonic fibroblasts and human cell lines were mined from public GEO database (Busslinger et al., 2017; Hamperl et al., 2017; Holzmann et al., 2019; Pope et al., 2014; Sanz et al., 2016) (**Tables S4 and S6**). The raw ChIP-seq sequence data were downloaded using Faster Download and Extract Reads in BAM format from NCBI SRA on Galaxy server (Afgan et al., 2018; Leinonen et al., 2011), and then were mapped against the human hg19 reference using bowtie2 (Langmead and Salzberg, 2012). For the biological replicate experiments, the resulting alignments from each experiments were combined using samtools merge (Danecek et al., 2021). Peaks were called by MACS2 with the parameters settings described previously (Busslinger et al., 2017; Feng et al., 2012). The BED file of called R-loop peaks of mouse embryonic fibroblasts was directly downloaded from GEO database (GSM1720621). The peak overlaps were determined using BEDTools suite with default parameters (Quinlan and Hall, 2010). The genomic annotation of peak overlaps was analyzed using ChIPseeker R package (Yu et al., 2015). The read density profiles and heatmaps of R-loop across overlapped regions were analyzed using deeptools (Ramírez et al., 2016). The significance of peak overlaps were calculated using bedtools fisher, ChIPseeker with nShuffle of 10000 and Genomic HyperBrowser with Monte Carlo sampling of 10000 (Sandve et al., 2010) (**Tables S5 and S7**).

**Hi-C analysis**

Hi-C data from MEFs were mapped at 1 kb resolution with the mm9 genome assembly and distiller pipeline (https://github.com/mirnylab/distiller-nf; version 0.0.3). The mapped data were converted to cooler files (Abdennur and Mirny, 2019) and balanced by iterative correction (Imakaev et al., 2012). Pile ups on R-loops were performed similarly to pile ups described previously (Banigan et al., 2022). Snippets of "observed" 5-kb-resolution Hi-C contact maps centered on R-loops, which were previously determined by DRIPc-seq (Sanz et al., 2016). Observed snippets were normalized by the contact probability scaling ("expected") to produce “observed-over-expected” snippets. Observed-over-expected snippets were averaged together to produce the average Hi-C contact enrichment maps. Gene annotations from GENCODE (https://www.gencodegenes.org/) were used to identify intra- and intergenic regions (Frankish et al., 2021).
